# *In situ* single-cell CRISPR screens uncover context-dependent rewiring of glioma states and immune recruitment

**DOI:** 10.64898/2026.11.08.765171

**Authors:** Nicholas Hou, Ioana-Teodora Lia, Ross M. Giglio, Bailey T. Van, Sebastian Pallais-Aks, Mathini Vaikunthan, Vivian Lu, Jonathan Guo, Destinee Williams, Tiffany C. Nieto, Lingting Shi, Cody Slater, Elham Azizi, José L. McFaline-Figueroa

## Abstract

*In vivo* single-cell CRISPR screens can resolve perturbation effects in native tissues, but few combine cell-type-restricted delivery, somatic tumor initiation, and combinatorial perturbation. Here we developed an RCAS-TVA-based single-cell CRISPR screening approach coupling oncogenic transformation with single- or dual-guide perturbation in defined neural progenitors. In glioma, tumor suppressor loss produced distinct genotype-dependent transcriptional states, including an *NF1*-loss-associated increase in mesenchymal programs. Introducing a fixed *Cdkn2a*(ARF) perturbastion altered malignant-cell state composition, immune-cell abundance, and the effects of additional tumor-suppressor perturbations, including increased representation of *Pten*- and *Rb1*-deficient cells, suggestive of the relief of an ARF-dependent genetic barrier. These perturbations converged on transcriptional states resembling pharmacologic p53 activation in human glioblastoma. Across mouse tumors and human glioblastoma models, this response was associated with increased astrocyte-like state representation. These findings establish a framework for single-cell analysis of context-dependent somatic perturbations during tumor initiation *in vivo*.

## Introduction

Pooled CRISPR screens have revolutionized functional genomics, enabling systematic, large-scale characterization of gene function. Coupling these screens with single-cell transcriptomics^1–4^ has extended this paradigm beyond fitness-targeted molecular readouts (proliferation, reporter expression)^5^. Perturb-seq and related approaches have enabled high-dimensional mapping of gene regulatory networks, cellular states, and responses to environmental or therapeutic perturbation^1,3,4,6–8^. More recently, these strategies have been extended *in vivo*^2,9–12^, allowing investigation of the effect of genetic perturbation in the context of native tissues and organs and defining how gene function varies due to complex cellular interactions.

Current *in vivo* single-cell CRISPR methods use delivery strategies that can make restriction to a defined cell population challenging (e.g., lentiviral delivery^9^, reliance on AAV tropism^13,14^, convection-enhanced delivery into established tumors^15^) or require additional regulatory or genetic systems to restrict perturbations to a defined population. Other approaches necessitate complex mouse engineering, *ex vivo* manipulation^2^, or early embryonic delivery of perturbations^9^, making them less suited to modeling somatically acquired mutations in a defined cell of origin. These limitations matter especially in cancer, where transformation arises from combinations of somatic alterations acquired in a cell of origin^16^, and where complex interactions between the tumor and the surrounding microenvironment shape the resulting phenotype.

Large-scale efforts have defined the genomic landscape of GBM, identifying near-ubiquitous overactivation of receptor tyrosine kinase (RTK) signaling (*EGFR*, *PDGFRA* activation), loss of negative regulation of the RAS/MAPK and PI3K effector pathways (*NF1*, *PTEN* loss), and inactivation of the *Rb* and *p53* tumor suppressor pathways (*CDKN2A/B, TP53* loss & *CDK4* activation)^17,18^. Bulk transcriptomic studies defined classical, proneural, and mesenchymal GBM subtypes^19^, while single-cell studies revealed substantial intra-tumor heterogeneity, hybrid programs, and a continuum of malignant cell states. These cell states are associated with recurrent drivers such as NF1 loss with MES-like programs and *CDK4* loss with NPC-like programs^20–22^. Moreover, these states exhibit high plasticity, shifting with treatment and at recurrence^23–25^. Studies of immune checkpoint blockade (ICB)-treated GBM show that baseline transcriptional state can predict survival and that subtype evolution under ICB is distinct from standard chemoradiation^26^. Perturbation screens that systematically vary genetic context during tumor initiation could complement these patient-derived datasets by defining how specific genetic architectures shape malignant-cell states and remodel the surrounding TME.

Here, we present Patchwork, an *in situ* single-cell CRISPR screening platform that adapts the **R**eplication-**C**ompetent **A**vian leukosis virus long terminal repeat with a **S**plice acceptor (**RCAS**) avian retroviral system for cell-type-specific delivery^27^ of single or combinatorial perturbations together with single-cell RNA-seq-compatible guide capture in cells expressing the avian viral receptor TVA under cell type or lineage specific promoters. RCAS-TVA has been used to model cancer in defined cells of origin, including in glioma, pancreatic, melanoma, ovarian, and breast cancer^27–29^, and has more recently been paired with CRISPR-Cas9 for somatic editing *in vivo*^30^. Patchwork is deployable in any tvA mouse line, delivering an oncogenic driver and one or more sgRNAs to a genetically defined population while resolving guide identity in every cell. We applied Patchwork in Cas9 Nestin-tvA mice to initiate gliomas by transforming neural progenitor cells in the subventricular zone and performed single- and dual-guide screens targeting canonical GBM tumor suppressors and exploratory loss-of-function events. Across these screens, Patchwork identified established and novel genotype-state relationships, revealed how *Cdkn2a*(ARF) loss reshapes perturbation-specific malignant programs and tumor-immune interactions, and uncovered a shared transcriptional response to *Pten* and *Rb1* loss that mirrors the response of human GBM to pharmacological p53 activation.

## Results

### Patchwork enables cell-type-specific single-cell CRISPR screens *in vivo*

Patchwork is an *in situ* single-cell CRISPR screening system built on the RCAS-TVA avian retroviral platform (**Figure 1a**). RCAS, an avian sarcoma/leukosis virus derived from the Rous sarcoma retrovirus, infects mammalian cells only when they express the cognate avian TVA receptor, allowing delivery of genetic payloads to genetically defined populations expressing *tvA* under a cell type-specific promoter. RCAS vectors accommodate transgenes on the order of the ∼2kb, similar to the size of the v-src coding sequence replaced from the Rous sarcoma retrovirus. We used this approximate limit to insert oncogenic transgenes with one or more CRISPR sgRNAs with varying sgRNA backbones. Prior work has shown that this approach enables somatic transformation and controlled genetic perturbation within a defined cell of origin, and has led to the development of multiple cell-type-specific tvA-expressing murine models^27,28^.

**Figure 1.**
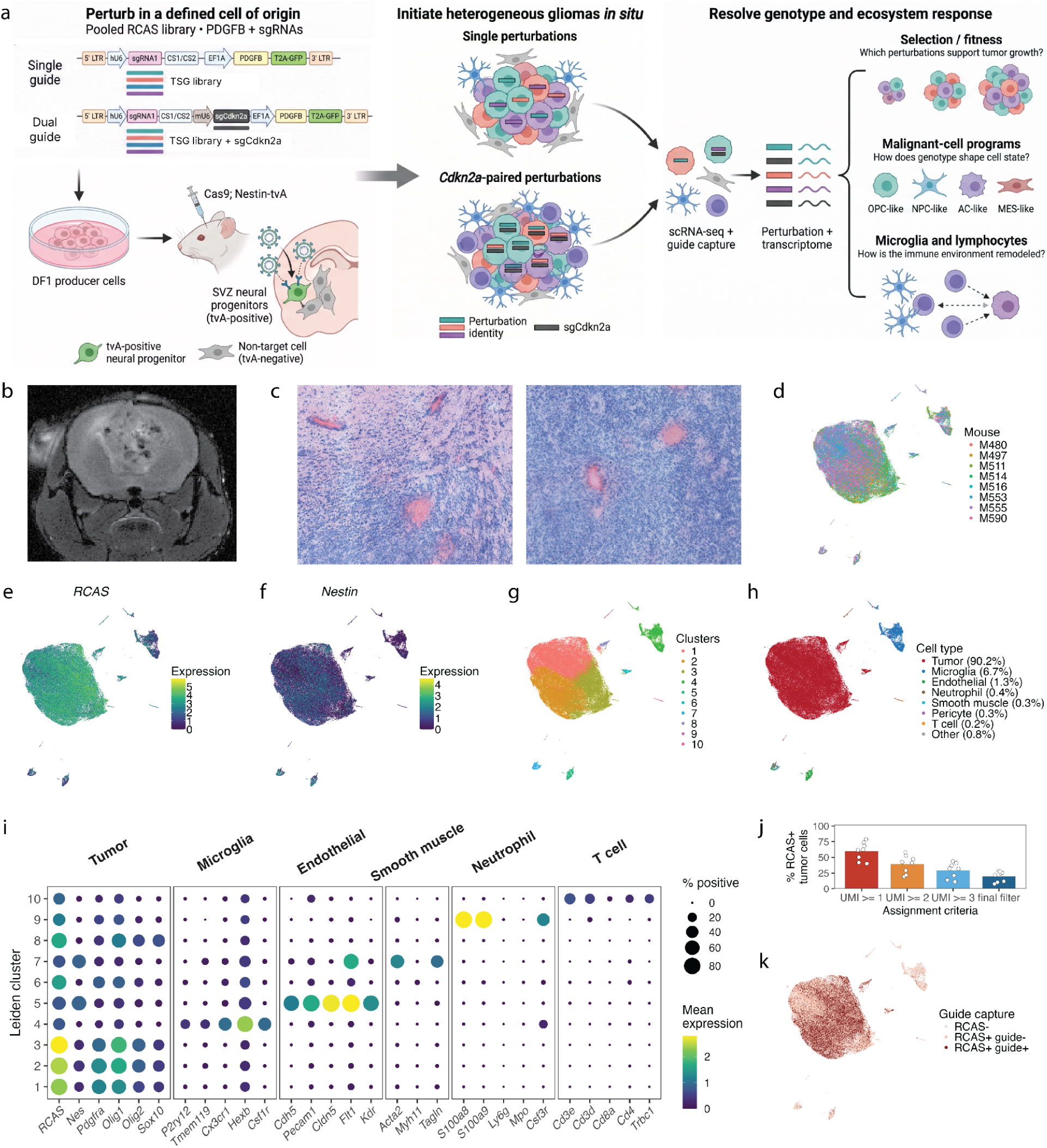
Patchwork enables cell-type-restricted somatic CRISPR screening during glioma initiation. **a,** Schematic of the Patchwork tumor-induction pipeline. DF1 producer cells are transfected with pooled RCAS-CS2-sg(TSG)-PDGFB libraries carrying guides against eight tumor suppressors (*Brd4*, *Cdkn2a*, *Cdkn2b*, *Cdkn2c*, *Nf1*, *Pten*, *Rb1*, *Trp53*), two exploratory loss-of-function targets (*Idh1* and *Idh2*), and a negative control (*Mdm2*), plus non-targeting controls (Ntc), in single-guide and *Cdkn2a*-paired dual-guide configurations. The transfected DF1 cells were injected intracranially into the subventricular zone of Rosa26-Cas9; Nestin-tvA mice, restricting transduction to Nestin⁺ neural progenitors. Created in BioRender. McFaline-Figueroa, J.L. (2026) https://BioRender.com/cpprxuc **b,** T2-weighted MRI of a Rosa26-Cas9; Nestin-tvA mouse 25 days after intracranial injection of 1 × 10⁵ DF1 cells transfected with the single-guide Patchwork construct. **c,** Hematoxylin and eosin staining of a representative single-screen tumor. **d-g,** Uniform Manifold Approximation and Projection (UMAP) embeddings of 121,514 cells from 8 mice in the single-screen, annotated by mouse-of-origin (**d**), RCAS transgene expression (**e**), *Nes* expression (**f**), and Leiden cluster (**g**). **h,** Per-cluster dot plot of selected canonical markers for six cell-type modules; dot size is the percentage of cells with non-zero expression and color is mean expression within that cluster. An expanded marker panel is shown in **Supplementary Fig. 5**. Expression in **e**, **f** and **h** is size-factor-normalized log expression. **i,** UMAP annotated by cell type; legend percentages give the fraction of all cells per type. **j,** Percentage of RCAS⁺ tumor cells per mouse carrying a detected guide, at total guide UMI ≥ 1, ≥ 2 and ≥ 3, and after the final guide-assignment filter (**Methods**). Dashed line, pooled rate across all eight tumors (total detected / total RCAS⁺ tumor cells). **k,** UMAP annotated by joint RCAS and guide detection: RCAS^−^, RCAS⁺/guide^−^, and RCAS⁺/guide⁺, showing guide capture relative to transduction efficiency. Guide detection is defined by the final assignment in (**j**).

We adapted the RCAS system for single-cell CRISPR screening by engineering RCAS vectors where perturbation identity can be recovered by direct guide capture using capture sequences compatible with commercial microfluidic technologies^1,31^. We generated a set of Patchwork vectors supporting single- or dual-guide delivery, a pol II open reading frame, and the varied sgRNA-capture configurations (**Supplementary Fig. 1a**). We performed *in vitro* validation to confirm TVA-dependent transduction, functional sgRNA delivery, and guide recovery by single-cell RNA sequencing (**Supplementary Fig. 1b-e)**. As expected, T98G dCas9-KRAB cells expressing TVA were transduced by RCAS vectors carrying sgRNAs targeting *HPRT1*, whereas TVA-negative cells were not (**Supplementary Fig. 2**). Under 6-thioguanine selection, sgRNAs targeting *HPRT1* conferred the expected resistance phenotype, confirming functional sgRNA delivery. Additional comparisons of two guide-capture configurations showed that CS1 provided higher guide recovery, whereas CS2 produced more uniform functional knockdown (**Supplementary Fig. 1e**). We therefore selected the CS2 configuration for *in vivo* perturbation experiments.

To determine whether Patchwork could be used to resolve the consequence of tumor induction given a variety of mechanisms, we designed a pooled CRISPR screen targeting GBM relevant tumor suppressors and pathway regulators (*Brd4*, *Cdkn2a*, *Cdkn2b*, *Cdkn2c*, *Idh1*, *Idh2*, *Nf1*, *Pten*, *Rb1*, and *Trp53*) together with negative (*Mdm2*) and non-targeting controls (*Ntc*) (**Fig. 1a)**. The Patchwork vector additionally encodes the activator of receptor tyrosine kinase signaling, *PDGFB*, under the control of a constitutive promoter. Including *Mdm2*, which is not expected to cooperate with *Pdgfb*-driven transformation, and Ntc, provides internal controls for tumors driven by *Pdgfb* overexpression alone, which can occur at low penetrance in the RCAS Nestin-tvA model^28^. Patchwork virus was generated in DF1 chicken cells, green fluorescent protein (GFP)-positive producer cells enriched by sorting with fluorescence-activated cell sorting (FACS), and delivered intracranially to the subventricular zone of adult Rosa26-Cas9 *Nestin*-tvA mice (**Supplementary Fig. 2-3**).

Tumors were harvested after confirmation by MRI as a discrete, clearly delineated mass ≥2 mm in diameter (**Fig. 1b**), or earlier if an animal showed signs of intracranial pathology, ≥20% weight loss, or declining body condition, per euthanasia criteria (**Methods**). Patchwork tumors displayed histological features of high-grade gliomas, including diffuse infiltration, extensive necrosis, and pseudopalisading necrosis (**Fig. 1c**). Following single-cell sequencing and quality control (**Methods**), we recovered 121,514 cells from 8 murine glioma tumors (**Fig. 1d**). Tumor cells expressed canonical glioma lineage markers (*Olig2*, *Gfap*) and were highly proliferative (*Mki67*), consistent with high-grade glioma (**Supplementary Fig. 4**). As expected, RCAS transgene and *Nestin* expression were highly enriched in tumor cells, which accounted for the majority (90.2%) of recovered cells (**Fig. 1d,e**). Microglia were the dominant non-malignant population (6.7%), followed by endothelial cells (1.3%) and neutrophils (0.4%), with smaller perivascular (smooth muscle 0.27%, pericyte 0.26%), monocyte-derived macrophage (0.19%) and T-cell (0.19%) compartments (**Fig. 1f-h**).

RCAS transgene was detected in 84.7% of recovered tumor cells, but guide capture was less efficient. Of RCAS-expressing cells, 60.5% carried at least one sgRNA UMI, with recovery varying from 40.1% to 78.8% across tumors (**Fig. 1i-j**). Recovery decreased monotonically with stricter thresholds, and 20.0% of RCAS-expressing cells were assigned a single perturbation under the strict filtering criteria (7.8% to 28.2% across tumors) (**Fig. 1j**). Guide recovery was therefore limited by the capture of any guide transcript as well as the assignment of captured guides to a single perturbation. Per-tumor recovery did not correlate with sequencing depth (**Supplementary Table 3**).

### Tumor-suppressor loss reshapes glioma cell state *in vivo*

Recovery of sgRNAs targeting tumor suppressor in our screen was largely consistent across individual mice (**Fig. 2a),** with sgRNAs targeting *Trp53* enriched in tumors relative to their representation in transfected DF1 cells (**Fig. 2b**; Fisher’s exact test, Padj < 10^-296^) and was the top or second most recovered genotype in all mice. *Nf1* sgRNAs were also enriched at the cohort level (Fisher’s exact test, Padj < 10⁻⁴⁷). As expected, Ntc and sgRNAs targeting *Mdm2* were amongst the least captured in our screen as were sgRNAs targeting *Pten* and *Rb1*. This relative enrichment of *Trp53* compared with *Pten* is consistent with prior RCAS-TVA studies showing stronger cooperation between *Trp53* loss and *PDGFB*-driven gliomagenesis, whereas the effect of *Pten* loss depends on age at tumor initiation with higher penetrance in pups compared to adult mice^28^.

**Fig. 2.**
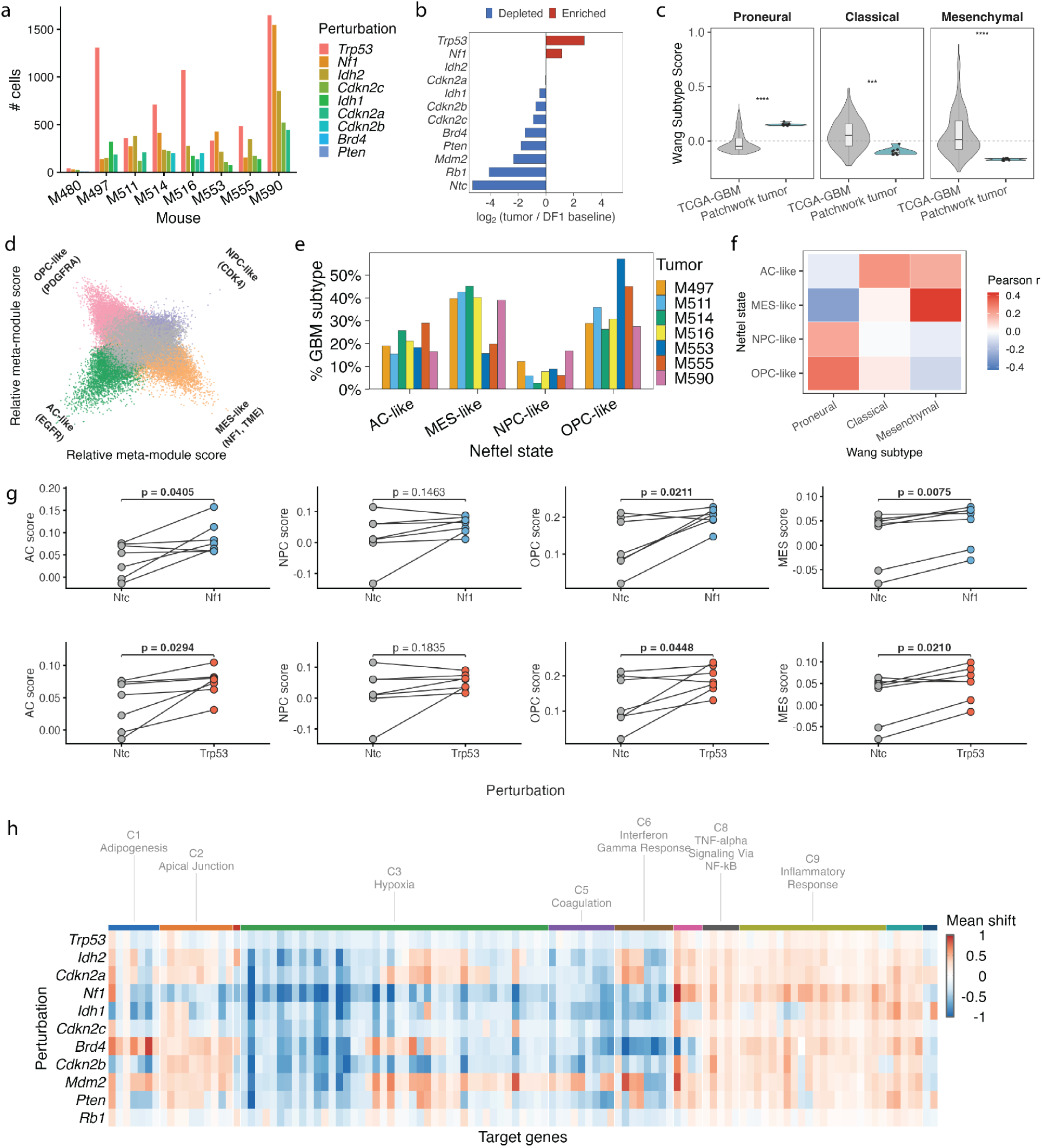
Tumor suppressor loss reshapes the cell-state landscape of Patchwork gliomas. **a,** Cells recovered per target gene in each single-screen tumor. The top five recovered gene targets per tumor are displayed. **b,** Enrichment or depletion of each target gene relative to the injected DF1 library, log₂(tumor/DF1). **c,** Wang 2017^20^ subtype scores in TCGA-GBM bulk tumors (n = 391) and single-screen Patchwork pseudobulk profiles (n = 8). Scaled expression scores of the aggregate expression of subtype signature genes relative to expression-matched control genes, zero (dashed line) denotes no enrichment over background. Boxes, median and IQR; whiskers, 1.5× IQR (***Padj < 10⁻³, ****Padj < 10⁻⁴, two-sided Wilcoxon rank-sum, Benjamini-Hochberg across subtypes). **d,** Neftel state representation^21,25^ of single-screen tumor cells annotated by state call. **e,** Neftel states fractions for the seven tumors with ≥ 50 resolved cells. **f,** Two-sided Pearson correlation between mean Neftel state score and mean Wang subtype score across all single-screen tumor cells. **g,** Per-tumor Neftel state scores in Ntc versus *Nf1* KO (top) or *Trp53* KO (bottom) cells. Scores are the log-transformed sum of size-factor-normalized counts over each signature (**Methods**). P-values are computed with two-sided Mann-Whitney U on all KO versus all Ntc cells pooled across tumors, unpaired and at the level of cells. **h,** Signed mean shift (relative to Ntc) per significant target gene as defined by SHERLOCK across perturbations (**Methods**). Columns are unsupervised gene clusters, each labeled with its top Hallmark term by GSEA over-representation analysis.

For our pilot single genotype screen, we first asked whether Patchwork tumors span the expression programs that define molecular subtypes of human GBM. We scored tumor cells for the proneural, classical, and mesenchymal programs defined by Verhaak and colleagues^19^. Patchwork tumors were biased toward the proneural axis relative to bulk expression profiles across the TCGA GBM cohort, with higher proneural and lower classical and mesenchymal scores (**Fig. 2c**), consistent with *Pdgfb*-driven tumor initiation in *Nestin*-expressing neural progenitors. Applying the subtype framework from Neftel et al. showed that Patchwork tumor cells occupy all four regions of the oligodendrocyte progenitor-, neural progenitor-, astrocyte-, and mesenchymal-like (OPC, NPC, AC, MES) cell state landscape (**Fig. 2d**). However, the OPC-like cell state was predominant in Patchwork tumors, consistent with the association of this state with alteration in *PDGFRA*. Overall, state composition was similar across tumors (**Fig. 2e**).

The Verhaak and Neftel frameworks captured related but non-identical features of Patchwork tumor-cell identity (**Fig. 2f**). OPC-like and NPC-like cells were positively associated with the proneural program, whereas MES-like cells were associated with the mesenchymal program. In contrast, AC-like state assignment showed little correspondence with the classical program, consistent with the relative absence of the EGFR-associated classical axis in our *Pdgfb*-driven model. Thus, Patchwork tumors span the major single-cell states observed in human GBM while retaining a pronounced OPC/proneural bias.

Examining subtype scores by genotype revealed that both *Nf1* and *Trp53* loss increased OPC-like and MES-like program scores relative to Ntc (**Fig. 2g**; all q < 0.05, Wilcoxon rank-sum with Benjamini-Hochberg correction). The mesenchymal shift after *Nf1* loss aligns with the strong association between NF1 inactivation and mesenchymal GBM in human tumors^19^ and with experimental studies linking NF1 loss to the induction of mesenchymal transcriptional programs^32–34^. No perturbation altered NPC-like programs, although *Cdkn2a* and *Cdkn2c* loss trended toward increased AC-like scores (FDR = 0.10 and 0.08, respectively).

Across the individual perturbations, we used a negative binomial generalized mixed effects model (GLMM) to identify differentially expressed genes (DEGs) relative to other tumor cells while controlling for tumor-specific confounders. Gene-level responses were modest, totaling 571 DEGs across 467 unique genes and ranging from 187 DEGs for *Trp53* knockout (KO) to 3 for *Pten* KO (median 61; FDR < 0.05, **Supplementary Fig. 7**). Notably, *Idh1* was itself induced among *Idh2* KO cells suggesting paralog compensation. Enrichment analysis on the per-perturbation DEGs found *Trp53* KO suppressing proliferative and oxidative metabolism (MYC targets NES = −2.74) while inducing hypoxia and cholesterol homeostasis. *Nf1* KO moved the same axes in the opposite direction (oxidative phosphorylation +1.69, MYC targets +1.68).

Modeling perturbations jointly with SHERLOCK²³, a structured variational autoencoder separating intrinsic from perturbation-specific variation, highlighted strong similarity between tumor cells across genotypes. Perturbations target gene weights clustered into programs common to most KOs, namely changes in, cholesterol homeostasis, oxidative phosphorylation, glycolysis and heme metabolism alongside IL-6/JAK/STAT3, interferon-γ and IL-2/STAT5 signaling (**Fig. 2h**). SHERLOCK therefore recovers a convergence on acquired programs across individual genotypes and tumors in our single-tumor suppressor loss of function screen.

### *Cdkn2a*(ARF) loss remodels the glioma microenvironment

To determine how genetic context alters the effect of tumor suppressor loss, we extended Patchwork to combinatorial perturbation using a dual-guide RCAS vector that delivers two sgRNAs together with *Pdgfb*. We fixed one sgRNA to target *Cdkn2a*, one of the most frequently altered tumor suppressors in GBM^17,18^, and paired it with the same perturbation library used in the single-screen. The fixed *Cdkn2a* guide edits within exon 1β, 87 nucleotides into the p19ARF open reading frame and is not expected to affect the reading frame encoding the cyclin-dependent kinase inhibitor p16INK4A. In RCAS/TVA models, loss of ARF substantially increases the rate of gliomagenesis compared to loss of Ink4a. In addition, targeting of ARF only allows us to interrogate the p16INK4A-Rb pathway within the same screen. Therefore, our ARF loss co-perturbation framework provides a minimal permissive genetic background that facilitates tumor establishment.

Using the same tumor-initiation and single-cell profiling workflow, we recovered 95,261 cells from ten mice. After quality control and integration, dual-screen cells resolved into ten Leiden clusters that mixed across the ten mice (**Fig. 3a**), with tumors forming a shared cell-state space. Similar to our single-genotype screens, dual-screen tumor cells spanned all four Neftel subtype states (**Fig. 3c**), but their distribution differed from the OPC-dominant single-screen tumors (**Fig. 3d**). The OPC-like compartment decreased from 46% to 18% of resolved tumor cells, while AC-like cells rose from 20% to 36%; NPC-like from 7% to 13% and MES-like from 27% to 33%. ARF co-perturbation therefore appears to reroute the malignant-cell landscape away from the OPC-dominant single-screen composition and toward AC-like, NPC-like and MES-like states.

**Figure 3.**
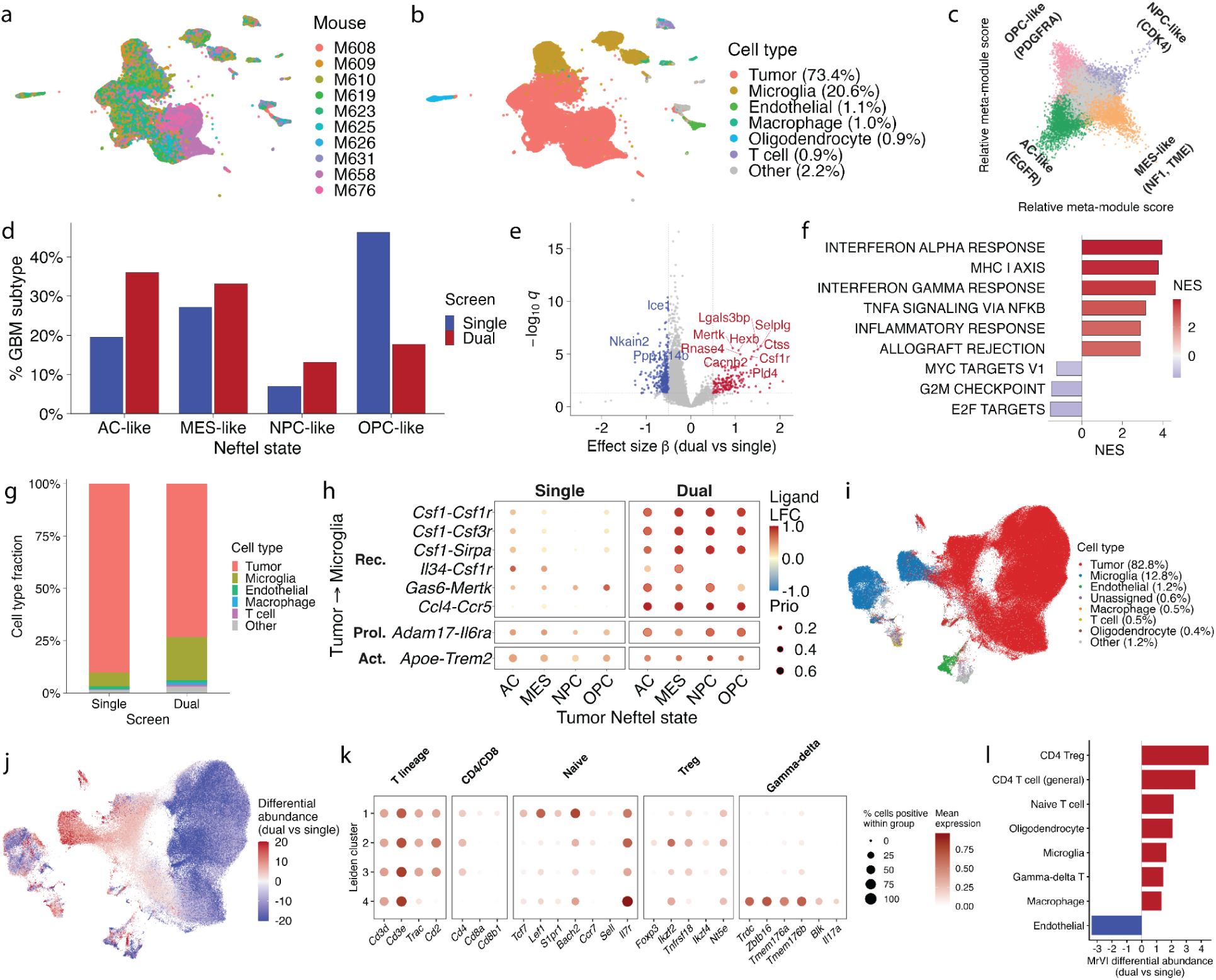
*Cdkn2a*(ARF) combinatorial perturbation reroutes the tumor subtype landscape and remodels the tumor immune microenvironment. **a**, UMAP of cells from our dual *Cdkn2a*(ARF) and tumor suppressor edited screen annotated by mouse of origin. **b**, Same as in (**a**) annotated by cell type. **c**, Neftel-state landscape of dual-screen tumor cells; cells colored by Neftel state call. Quadrants are annotated with the canonical driver corresponding to each state. **d**, Neftel states fractions in the single versus dual screens, across all tumor cells. **e**, Volcano plot of the results of differential expression analysis between dual- vs single-screen tumor cells. A negative-binomial GLMM with mouse as a random effect was applied. Dashed lines mark absolute effect size |β| = 0.5 and q-value = 0.05. **f,** GSEA of DEGs from (**e**) using the MSigDB Hallmarks gene set collection^41^. Bars are colored by NES; FDR < 0.05. **g**, Cell-type fractions in the single-versus dual-screens. **h**, MultiNicheNet tumor-to-microglia signaling by tumor Neftel state and screen; dot size corresponds to prioritization score, and color scales with log fold change. **i,** Joint single- and dual-screen UMAP of the MrVI sample-unaware (u) latent space, annotated by cell type; legend percentages give the fraction of total cells. **j**, Same UMAP annotated by MrVI differential abundance (dual versus single log ratio). **k**, T cell subset annotation by Leiden sub-clustering of the T cell-assigned clusters across both the single- and dual-screens. **l**, MrVI differential abundance by TME cell type.

We first inspected the transcriptional consequences of *Cdkn2a*(ARF) co-perturbation across all tumor cells. We performed differential expression using a negative-binomial mixed-effects model and gene-set enrichment analysis (GSEA) (**Fig. 3e,f**). Dual-screen cells strongly upregulated interferon-γ response, interferon-α response, TNFα-NFκB signaling, inflammatory response and MHC class I antigen presentation, while cell-cycle programs (E2F targets, G2M checkpoint, MYC targets V1) were suppressed relative to single-screen cells. Up-regulated DEG genes included factors associated with tumor-intrinsic IFN-γ/MHC-I induction (*Cst3*, *Lpcat2*), microglia-identity gene expansion (*Csf1r*, *Selplg*, *Olfml3*, *Hexb*, *Ctss*), and presence of parenchymal oligodendrocyte (*Mbp*, *Mobp*). The dual screen thus exhibits an increased inflammatory phenotype associated with tumor-cell state remodeling and shifts in the tumor microenvironment (TME) composition.

Comparing non-tumor compositions between screens, the immune compartment expanded substantially under combinatorial *Cdkn2a*(ARF) loss, particularly microglia which increased approximately three-fold (6.7% of single-screen cells to 20.6% of dual-screen cells) (**Fig. 3g**). To dissect tumor-to-microglia signaling at the ligand-receptor resolution, we performed MultiNicheNet analysis on microglia receivers with the tumor senders stratified by Neftel state and screen (**Fig. 3h**). *Cdkn2a*(ARF)-loss resulted in four distinct recruitment axes shared across all four Neftel states, the CSF1 multi-receptor hub (*Csf1*-*Csf1r*/*Csf3r*/*Sirpa*), *Ccl4*-*Ccr5* chemokine recruitment, *Gas6*-*Mertk* TAM-receptor engagement, and *Adam17*-*Il6ra IL*-*6* trans-signaling. Both *CSF1R* microglia ligands were enriched in the dual-screen with inflammatory *CSF1* far more so than the homeostatic *IL-34* ligand, suggesting that the dual-screen tumor compartment engages *CSF1R* predominantly through the inflammatory ligand.

Jointly embedding tumor and TME cells across both screens using Multi-resolution Variational Inference (MrVI)^35^, a deep generative model that learns a cell-state representation invariant to sample-of-origin, recovered a shared cell-type architecture spanning tumor, microglia, macrophage, T cell, endothelial and oligodendrocyte populations (**Fig. 3i**). Per-cell differential abundance on the tumor effect-invariant latent embedding concentrated dual-screen enrichment in the T cell compartment, with microglia, macrophage and oligodendrocyte populations also dual-associated (**Fig. 3j**). Re-embedding and Leiden clustering the joint T cell compartment resolved four T cell subsets: naive T cell; Treg; γδ T cell; and an unpolarized CD4 T cell population expressing T-lineage markers without a dominant state program. Labels were validated with canonical lineage markers (**Fig. 3k**); Treg identity was assigned based on *Ikzf2*, *Tnfrsf18* and *Nt5e* alongside sparsely captured *Foxp3*. Each of these states has a counterpart in human GBM^36–39^, with the exception of the γδ IL-17 program in mouse tumors^40^. MrVI per-cell differential abundance ranked Tregs as the most dual-enriched population (**Fig. 3l**), rising from 19.6% of the T cell compartment in the single-screen to 29.2% in the dual.

Together, these findings suggest that constitutive *Cdkn2a*(ARF) loss remodels the TME with evidence for differential tumor-intrinsic engagement of an interferon-γ / antigen-presentation programs (**Fig. 3e**), tumor-microglia recruitment (**Fig. 3h**), and T cell reprogramming toward a Treg-enriched state (**Fig. 3j-l**).

### *Cdkn2a*(ARF) loss reshapes the molecular consequences of tumor-suppressor perturbation

Comparing guide representation between the single- and dual-guide screens revealed context-dependent selection for tumor suppressor loss (**Fig. 2b**, **4a**). Although *Trp53* and *Nf1* perturbation remained among enriched genotypes, *Cdkn2a*(ARF) loss increased the representation of *Pten*, *Rb1,* and *Cdkn2b*.

**Figure 4.**
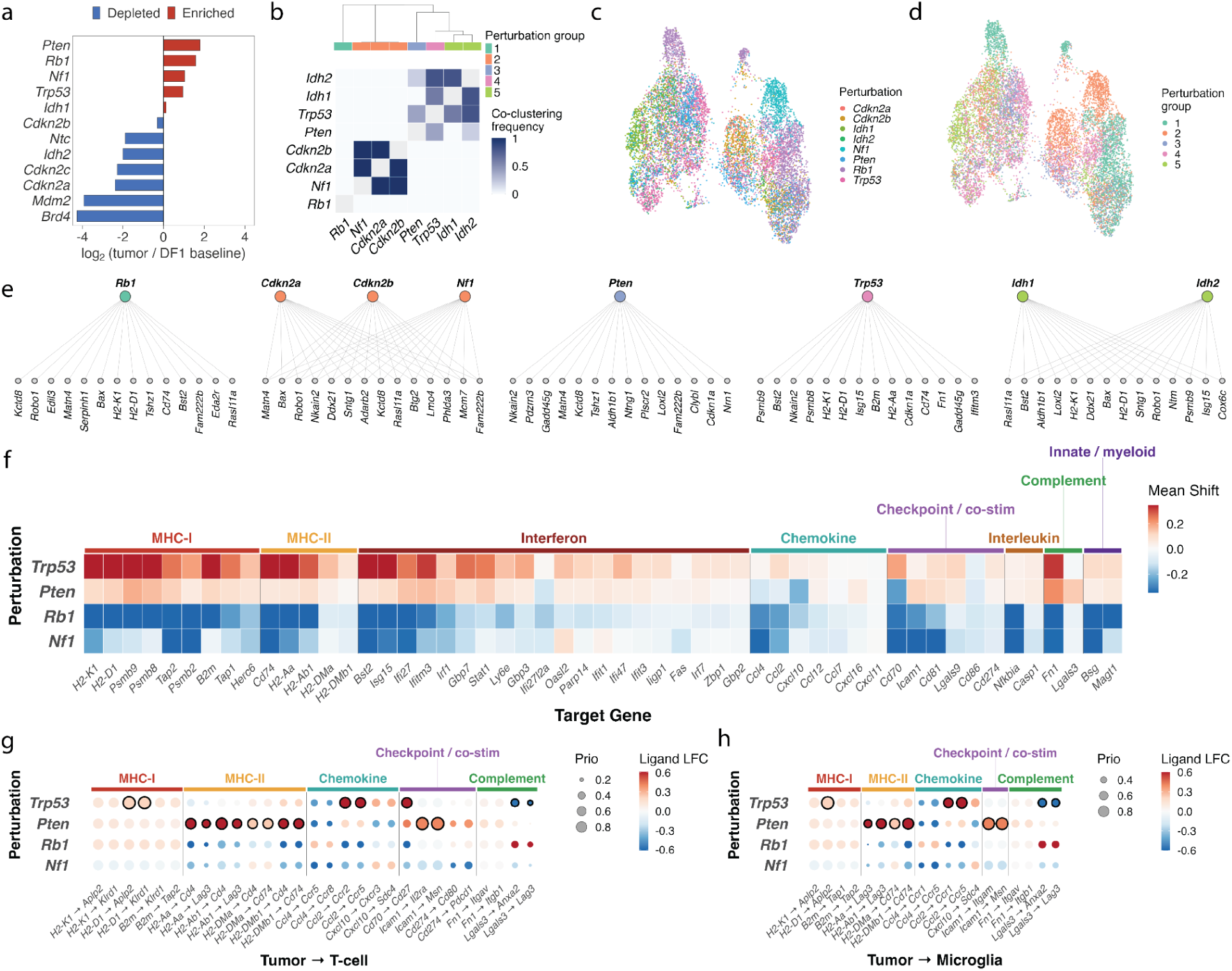
Genetic context determines perturbation-specific transcriptional programs *in vivo*. **a**, Enrichment or depletion of perturbations in dual-screen tumors relative to sgRNA frequency in the injected DF1 library. The non-targeting control (Ntc) is plotted alongside the targeted perturbations for reference. *Mdm2* depletion is the expected consequence of MDM2 loss in a *TP53*-wild-type, *ARF*-null background. **b,** Consensus co-clustering of perturbations across five SHERLOCK seeds. Within each seed, perturbations are grouped by hierarchical clustering of the SHERLOCK-inferred perturbation covariance. Each plot cell gives the fraction of seeds in which a pair falls in the same group. **c**, UMAP of the SHERLOCK latent *z* space annotated by perturbation. **d**, The same embedding annotated by consensus perturbation group. **e**, Significant target genes shared within each consensus perturbation group, as a bipartite graph. **f**, Signed mean shift in significant target genes relative to Ntc (**Methods**) for the top four enriched perturbations in the dual screen: *Trp53*, *Pten*, *Rb1* and *Nf1*. Genes in the x-axis denote immune-associated genes with a significant target gene across one or more perturbations. **g**, Tumor cells signaling to T cells computed with MultiNicheNet using mouse as the unit of replication. Ligand differential expression was tested by edgeR pseudobulk analysis of tumor-cell profiles, contrasting each genotype against the pooled remaining genotypes. Receptor expression was required in the receiving compartment. Fill gives the tumor-side ligand log₂ fold change, dot area the MultiNicheNet prioritization score, and a black ring an uncorrected P < 0.05 for that ligand. **h,** Same as (**g**) but for signaling to microglia.

Per-tumor distributions of the top five perturbations across the ten dual-screen tumors (**Supplementary Fig. 17a**) showed variable but broadly representative sampling of the perturbation panel across mice. Perturbations with cohort-total cell counts exceeding the Ntc control (n = 239 cells) were retained for downstream per-perturbation analyses (**Supplementary Fig. 12a**). Interestingly, guide-level composition differed between screens. Single screen tumors were typically dominated by one gRNA with the same guide prevailing in every mouse (**Supplementary Fig. 6c**) whereas dual screen tumors retained a more balanced within-tumor representation and greater between-tumor variability (mean pairwise Bray-Curtis 0.52 vs. 0.18; paired Wilcoxon signed-rank P = 0.031; **Supplementary Fig. 12c**). This supports our hypothesis that fixed *Cdkn2a*(ARF) loss provides a more permissive context for tumor evolution.

We next used SHERLOCK to more broadly characterize the shared regulatory structure across combinatorial perturbations. Consensus clustering of the learned perturbation covariance structure (**Methods**) resolved five groups across the eight retained perturbations (**Fig. 4b-d**): *Cdkn2a* with *Cdkn2b* and *Nf1*; *Idh1* with *Idh2*; and *Pten*, *Rb1* and *Trp53* each as singletons. The pairing of *Idh1* with *Idh2* is consistent with the two paralogs catalyzing the same NADP⁺-dependent reaction in different subcellular compartments^42–44^. The grouping of *Cdkn2a* with *Cdkn2b* is consistent with their adjacent genomic location and shared function as the CDK4/6 inhibitors p16INK4A and p15INK4B^45,46^.

To investigate perturbation-specific effects shared within each group, we analyzed target genes which are defined by SHERLOCK as the counterfactual effect of a given perturbation. Significant target genes shared within perturbation groups (**Fig. 4e**) comprised changes in: antigen-presentation and interferon-stimulated gene expression (*H2-K1*, *H2-D1*, *B2m*, *Cd74*, *H2-Aa*, *Psmb8*, *Psmb9*, *Bst2*, *Isg15*, *Ifitm3*); p53-pathway and cell-cycle effectors (*Bax*, *Cdkn1a*, *Gadd45g*, *Btg2*, *Phlda3*, *Mcm7*); and neural adhesion and matrix genes (*Robo1*, *Ntng1*, *Ntm*, *Nrn1*, *Nkain2*, *Matn4*, *Serpinh1*, *Loxl2*, *Fn1*). The antigen-presentation and interferon module was concentrated in the *Trp53* and *Idh* groups, while the *Cdkn2a*/*Cdkn2b*/*Nf1* group shared predominantly proliferative and p53-pathway targets.

Having demonstrated that constitutive *Cdkn2a*(ARF) loss remodels the immune compartment in Patchwork tumors, we next asked how individual genotypes contribute to this phenotype shift. The union of significant target genes across perturbations were grouped into co-regulated clusters (**Supplementary Fig. 17d**; **Methods**). One cluster consisted largely of interferon-stimulated genes. We classified genes in this cluster by immune function (antigen presentation, interferon response, chemokine, costimulatory) and examined the mean normalized target gene shift for the four perturbations enriched in dual tumors: *Trp53*, *Pten*, *Rb1* and *Nf1* (**Fig. 4f**). *Trp53* loss markedly raised MHC-I genes (*H2-K1*, *H2-D1*, *B2m*, *Tap1*, *Psmb8*, *Psmb9*). MultiNicheNet^47^ independently nominated *Trp53*-perturbed tumor cells as the strongest predicted chemokine senders to T cells and microglia (*Ccl2*-*Ccr2*, *Ccl2*-*Ccr5*) and as the strongest source of the costimulatory ligand *Cd70* (**Fig. 4g,h**). In our *Pdgfb*-driven Patchwork model p53 loss appears immune-engaging rather than immune-evasive. Interestingly, p53-chemokine crosstalk has also been reported in both glioblastoma^48^ and pancreatic cancer^49^. *Pten* loss produced a similar but weaker upward shift in the immune programs. MultiNicheNet nominated *Pten*-perturbed cells as the strongest predicted source of MHC-II signaling to both T cells and microglia (**Fig. 4g,h**). *Rb1* and *Nf1* loss each resulted in negative or relatively muted perturbation effects across the immune function modules.

### *Pten* and *Rb1* loss converge on p53 activation and AC-like persistence

Our results suggest that ARF co-perturbation altered the selection and molecular state of tumor-suppressor genotypes, in particular, an enrichment for *Pten* and *Rb1* loss in our combinatorial screen (**Fig. 4a**). We therefore asked what transcriptional programs accompanied their increased tolerance in this genetic background. *Pten*-deficient cells showed significant activation of p53 transcription and oxidative phosphorylation and suppression of G2M, E2F and MYC programs suggestive of a p53-high, proliferation-low state (**Fig. 5a**).

**Figure 5.**
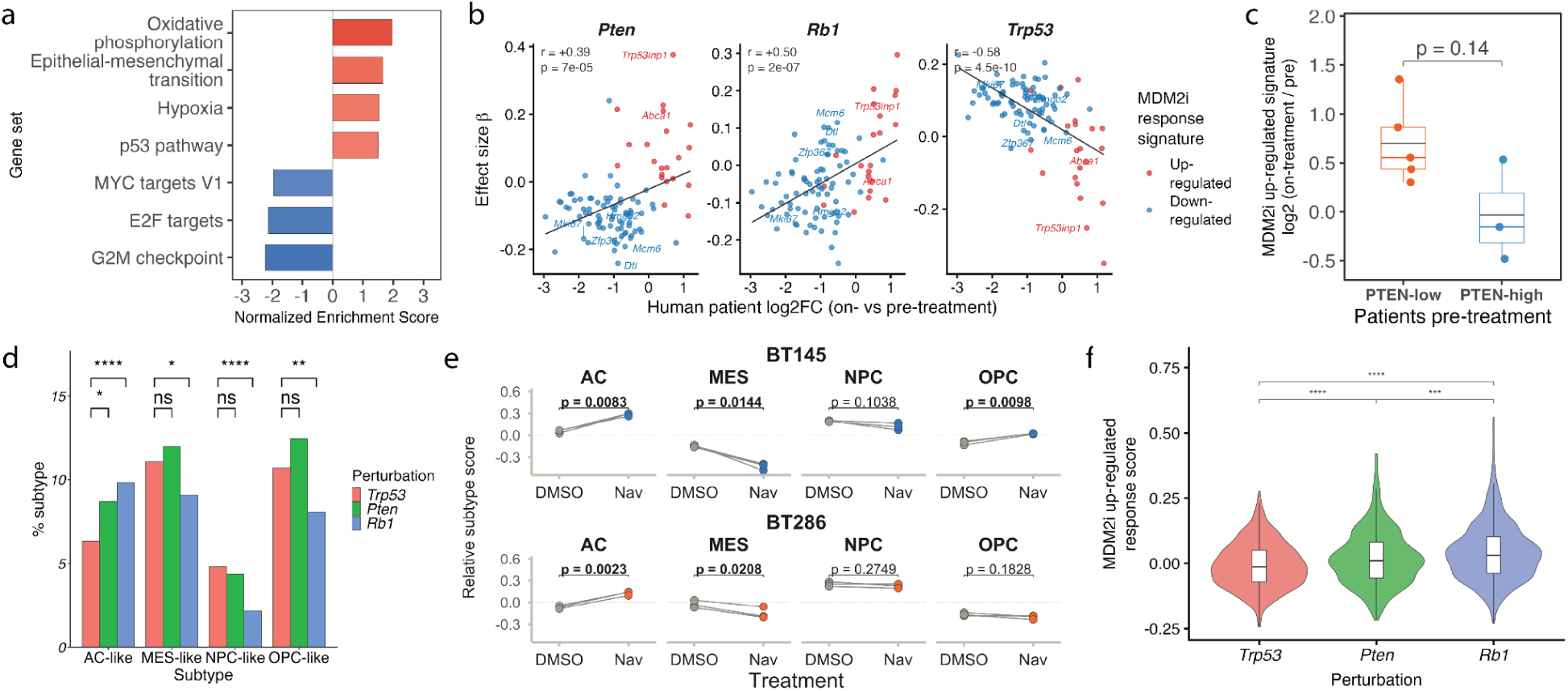
*Pten* and *Rb1* loss phenocopy pharmacologic p53 activation and associated AC-like persistence. **a**, GSEA of *Pten* KO dual-screen differential expression against the Hallmark collection. Bars are normalized enrichment scores, all shown at Padj < 0.05. **b,** Patchwork dual-screen KO βs correlation against mean human patient log₂ fold-change (navtemadlin-treated versus pre-treatment) across 96 orthologous navtemadlin-signature genes. Genes are annotated by the navtemadlin MDM2i response signature arm: up- or down-regulated. Pearson *r* and *p* are inset per plot. **c**, MDM2i up-regulated response signature, log₂(30 hours navtemadlin-treatment / pre-treatment) in the same eight patients. Patients are split at pre-treatment according to bimodal PTEN expression (**Supplementary Fig. 18c**). Exact Wilcoxon rank-sum P-value is reported by the bracket. **d,** Subtype fractions bar plots across *Trp53*, *Pten* and *Rb1* KOs in the dual screen. Brackets give the AC-like fraction of each KO against the *Trp53* reference by 2×2 chi-squared with continuity correction. Benjamini-Hochberg-adjusted is applied across KOs and subtypes; *Padj < 0.05, ****Padj < 0.0001. **e,** Neftel state scores shift across human GBM patient-derived neurospheres (navtemadlin-treated versus DMSO). The scores are centered within each cell line. Paired t-test, raw *P*-value inset per subtype. **f,** MDM2i up-regulated response signature score in *Trp53, Pten* and *Rb1* KO cells in the dual-screen. Two-sided Wilcoxon rank-sum, Benjamini–Hochberg-adjusted across the three; ***Padj < 0.001, ****Padj < 0.0001.

We first asked whether this response was similar to physiological p53 hyperactivation states in human GBM. Navtemadlin, an inhibitor of the E3 ubiquitin ligase MDM2, leads to the activation of wild-type p53 and has been evaluated in a window-of-opportunity trial in recurrent TP53 wild-type GBM^50^. The molecular effects of *Pten* loss in Patchwork tumors significantly correlated with the response to MDM2 inhibition (MDM2i) in paired biopsies pre- and post-navtemadlin treatment (**Fig. 5b**). *Rb1* KO cells also showed concordance with MDM2i response. Therefore, *Pten* and *Rb1* tumor suppressor perturbations with differential selection due to ARF disruption converge on a p53-active state.

*Pten* loss can activate mTORC1/2 signaling which can promote p53 stabilization^51–53^. Consistent with this possibility, the navtemadlin response was greater in PTEN-low versus PTEN-high patient tumors although the results were not significant across this limited patient cohort (**Fig. 5c**). RB1 loss disrupts the regulation of E2F activity, which can induce the ARF-MDM2-p53 checkpoint^54–58^. Constitutive ARF disruption could therefore attenuate stress response to RB1 loss. Consistent with this model, *Rb1* KO cells retained a positive p53-transcriptional response. Therefore, our results suggest that ARF constitutes a genetic barrier that limits the persistence of *Pten* and *Rb1* null cells with elevated p53 activity.

We next asked whether this p53-associated response was associated with specific tumor cell state change. Rendo et al. observed an enrichment for the AC-like signature in on-navtemadlin patient-biopsies. Our reanalysis of bulk RNA-seq data of navtemadlin-exposed BT145 and BT286 GBM neurosphere cultures also identified an MDM2i-dependent increase in the AC-like program and a decrease in MES-like program scores^50^ (**Fig. 5d**). Compared to p53 null cells, *Pten* and *Rb1* KO cells displayed an increase in the fraction of AC-like cells, suggesting that the AC-like state might be compatible with increased p53 activity (**Fig. 5e**).

Together, these findings connect *Pten* and *Rb1* loss, which become selectively enriched after ARF disruption, to transcriptional states that resemble p53 activation in human GBMs. In Patchwork tumors, AC-like cells persist despite engagement of this response (**Fig. 5f**), supporting a model for tolerance in which this state is compatible with sustained p53 activity. *PTEN* and *CDKN2A* loss frequently co-exists in high grade glioma and several studies suggest that *CDKN2A* loss precedes or accompanies *PTEN* loss during malignant transformation. Thus the permissive effect of ARF disruption on *Pten-*deficient cells identified by our *in vivo* screens is consistent with a recurrent evolutionary trajectory observed in human glioma.

## Discussion

We developed Patchwork, an *in situ* single-cell CRISPR screening platform that couples cell-type-restricted somatic perturbation to single-cell transcriptomic readout. High-complexity perturbation screens are readily implemented in transplantable tumor models, where cells can be engineered and expanded *ex vivo* before implantation^11,15,59^; Patchwork instead introduces pooled perturbations directly into a genetically defined endogenous cell population during somatic tumor initiation, preserving the native sequence of transformation, clonal selection and microenvironmental co-evolution. Importantly, our approach can be translated to existing tvA lines with minimal mouse engineering, and perturbations are introduced somatically allowing for the modeling of tumors at various stages in development (i.e., neonates, pups, adults). The modular vector series supports single- or dual-guide delivery alongside an oncogenic driver, so genetic background, driver, and perturbation can be varied independently within one experiment. Future work could also leverage split-vector or self-inactivating RCAS designs^60^ to increase packaging capacity, potentially enabling direct incorporation of Cas enzymes into the retroviral vectors.

Applying Patchwork to *Pdgfb*-driven gliomagenesis in Nestin-tvA mice recovered genotype-state relationships inferred from human tumors. *Nf1* loss shifted malignant cells toward the mesenchymal program, recapitulating the association between NF1 inactivation and mesenchymal GBM^19,21^; *Trp53* and *Nf1* guides were selected for during tumor formation.

Our results suggest that genetic context, not perturbation identity alone, determines the consequence of tumor suppressor loss. Fixing *Cdkn2a*(ARF) as a constitutive second perturbation altered which guides were selected (most strikingly enrichment of *Pten* and *Rb1*); rerouted the malignant-cell landscape from OPC-dominant toward coordinated AC-like and MES-like expansion; and produced CRISPR KO transcriptional responses uncorrelated with the same KO response in a *Cdkn2a*(ARF)-intact background. This context dependence cannot be readily captured from patient cohorts. ARF loss also remodeled the microenvironment including: changes in tumor-intrinsic interferon-γ and antigen-presentation programs; expression of genes related to tumor-to-microglia recruitment; increased expression of inflammatory *CSF1* relative to homeostatic *IL34* at the shared CSF1R receptor; and T cell expansion toward a Treg-enriched state.

Investigating the correspondence between our model and human GBM we find that *Pten* and *Rb1* KO responses in our dual screen correlate to the response of human GBM to the p53 activator navtemadlin. In the screen and in navtemadlin-treated human cells, we observe persistence of AC-like cells under sustained p53 activation. This result may explain clinical phenotypes including that navtemadlin-resistant tumors do not appear to carry TP53-inactivating mutations, and may highlight a non-genetic mechanism of resistance.

The Patchwork system provides additional opportunities for optimization. Although transduction of our target library does not appear to be limiting guide capture rate remains an efficiency-limiting step. While ∼85% of recovered cells expressed the RCAS transgene and ∼60% expressed a guide RNA, we confidently assigned 20% to a single perturbation. This constrains the number of perturbations that can be powered in one animal and sets the ceiling on library size. However, our results suggest that even our current designs can scale to hundreds of perturbations across reasonable mouse cohort sizes (**Supplementary Fig. 13**). Our conclusions also rely on a single cell of origin and a single oncogenic driver. *Pdgfb*-driven initiation from Nestin+ progenitors biases tumors toward proneural and OPC-like programs and largely omits the EGFR-associated classical axis, so the state landscape sampled here is a subset of human GBM rather than a representation of it. However, Patchwork vectors could be extended to model EGFR and CDK4 overactivation to model AC-like and NPC-like biased tumors and could incorporate *Nf1* loss in the absence of PDGF overexpression to bias toward a MES-like program^61^.

## Methods

### Plasmid construct design and cloning

We constructed four core Patchwork plasmids on the RCAS-BP-Y parental backbone, all sharing an EF1A-driven GFP reporter. The pilot construct (RCAS CS1/2 sgRNA) carries a single human U6 (hU6)-driven sgRNA cassette between the RCAS LTRs, followed by one of two direct-capture sequences (CS1 or CS2) compatible with 10x Genomics Chromium Single Cell 3′ Gene Expression with Feature Barcoding. The second construct adds *Pdgfb* downstream of the sgRNA cassette. The third adds a second sgRNA cassette driven by the mouse U6 (mU6) promoter with a distinct backbone, enabling dual-guide perturbation.

Variable sgRNAs were inserted at the sgRNA1 position by BsmBI-v2 (NEB) digestion and Gibson assembly (NEBuilder HiFi, NEB) of annealed oligonucleotides. The sg*Cdkn2a* cassette and the EF1A-PDGFB-T2A-GFP cassette were introduced by PCR amplification (NEBNext High-Fidelity, NEB) followed by Gibson assembly into the linearized backbone. Cloning was performed in a pDONR Gateway entry vector (Thermo Fisher) and recombined into RCAS-BP-Y with Gateway LR Clonase II to yield the replication-competent construct. All constructs were verified by whole-plasmid nanopore sequencing (Plasmidsaurus) and propagated in NEB Stable competent *E. coli*. Constructs will be deposited to Addgene.

### sgRNA design

Library sgRNAs were designed with CRISPick using Mouse GRCm38 (NCBI RefSeq v.108.20200622), CRISPRko mechanism and SpyoCas9 (NGG). The five highest-scoring protospacers per gene were selected for each of the 11 tumor suppressors, together with non-targeting controls. Protospacer sequences, target genes, on-target efficacy and aggregate off-target scores are found in **Supplementary Table 1**. Perturbations are referred to throughout by the gene each protospacer targets.

The fixed second guide of the dual-guide construct (5′-TGGTGAAGTTCGTGCGATCC-3′) is expressed from the mU6 promoter and carries no capture sequence, so it is absent from the feature reference and is never recovered in per-cell guide calls. Its genomic target was determined by BLASTN against the NCBI nucleotide collection. Every alignment at 100% identity and 100% query coverage was to *Cdkn2a* exon 1β or ARF records with no alignment to an exon-1α-only record. The cut site lies 87 nucleotides into the p19ARF open reading frame, upstream of the exon 2 sequence shared between reading frames, so the guide disrupts ARF while leaving p16INK4a intact.

### Cell culture

Avian DF1 producer cells (ATCC CRL-12203) and human glioblastoma T98G cells (ATCC CRL-1690) were obtained from ATCC. T98G cells stably expressing dCas9-BFP-KRAB were generated by lentiviral transduction with pHR-SFFV-dCas9-BFP-KRAB (Addgene 46911) and sorted by fluorescence-activated cell sorting (FACS) for high BFP expression^62^. A tvA-mCherry receptor line was generated by AAV transduction of a TVA-mCherry fusion using pAAV-EF1a-FLEX-TVA-mCherry (Addgene 38044). To ensure the TVA-mCherry is in frame, pAAV-EF1a-FLEX-TVA-mCherry was incubated with CRE in vitro prior to AAV generation. All lines were maintained in DMEM with 10% FBS and 1% penicillin-streptomycin at 37 °C, 5% CO₂.

### Viral production and transduction

DF1 cells were transfected with each RCAS construct using FuGENE HD (Promega) at a 1:4 DNA-to-reagent ratio, expanded to 10 cm dishes, and sorted by FACS for GFP on a BD FACSAria III. The same sorted preparations provided both virus-containing conditioned medium for in vitro transduction and the injection material for intracranial delivery.

For transduction, 150,000 T98G dCas9-KRAB cells per well were exposed to 3 mL of conditioned medium and spinfected^63^ at 800 × g for 60 min at room temperature with no enhancer. GFP was assessed at 96 hours by brightfield, GFP and dsRed microscopy (10X magnification, Zeiss Axio Observer). Functional CRISPRi delivery was assayed by treating transduced cells expressing CS1 or CS2 sgHPRT1 or sgNtc with 10 µM 6-thioguanine (Sigma-Aldrich A4882) for 96 hours, with relative growth quantified by manual counting and normalized to untreated controls.

### *In vitro* single-cell RNA-seq library preparation

6-TG-treated and untreated T98G dCas9-KRAB +TVA-mCherry cells transduced with RCAS-CS1/2-sgHPRT1 were dissociated with TrypLE Express and loaded into a 10x Genomics Next GEM Chromium Single Cell 3′ v3.1 kit with Feature Barcoding at a target of 3,000 cells per channel. Libraries were sequenced on an Illumina NovaSeq.

### Animal model and intracranial injection

All procedures were performed in accordance with protocols approved by the Columbia University Institutional Animal Care and Use Committee (IACUC) and conducted per NIH guidelines. Rosa26-Cas9 heterozygous;Nestin-tvA hemizygous mice were purchased from The Jackson Laboratories, maintained and crossed to generate Cas9 homozygous;Nestin-tvA mice. Mice of both sexes were used.

Adult mice (≥8 weeks) were anesthetized with isoflurane, immobilized in a stereotactic frame and given pre-emptive subcutaneous buprenorphine XR (3.25 mg/kg). FACS-sorted GFP⁺ DF1 cells were resuspended in 1 µL PBS and injected at 1.0 µL/min with a Hamilton 75N syringe into the subventricular zone (bregma 0 mm AP, −0.5 mm lateral, −1.5 mm from dura): approximately 100,000 cells per mouse for the single screen and 500,000 for the dual screen. Because the dual-guide construct exhibited lower transduction efficiency than the single-guide vector *in vitro*, the dual-screen DF1 dose was increased fivefold to approximately compensate for reduced effective viral delivery (**Supplementary Fig. 2c**). The needle was held for ∼1 min before withdrawal, the site sealed with bone wax and the scalp closed with Vetbond (3M). Mice were monitored daily for the first week and twice weekly thereafter for tumor progression and clinical signs (macrocephaly, lethargy, dehydration, hunched posture, ruffled coat, impaired locomotion, or ≥20% weight loss).

### Tumor monitoring by MRI

Tumor formation was assessed at 20 days post-injection by T2-weighted MRI on a Bruker BioSpec 9.4 T small-animal scanner using the Columbia University Irving Medical Center Oncology Precision Therapeutic and Imaging Core’s standard T2-weighted protocol. Mice were anesthetized with isoflurane and maintained prone on a heated support. Tumor presence was assessed visually; tumor volumes were not quantified.

### Tumor harvest, dissociation, and single-cell RNA-seq library preparation

Tumor-bearing mice (n = 8 single screen; n = 10 dual screen) were euthanized at endpoint criteria and tumors micro-dissected. Dissociation followed Slyper et al. (2020) using the Adult Brain Dissociation Kit (Miltenyi 130-095-942)^64^. Suspensions were filtered at 100 µm, washed in cold PBS + 1% BSA, resuspended in PBS + 0.4% BSA and loaded into a 10x Genomics Next GEM Chromium Single Cell 3′ v3.1 kit with Feature Barcoding at a target of 10,000 cells per channel. Gene expression and Feature Barcode libraries were sequenced on an Illumina NovaSeq. Per-library metrics are in **Supplementary Table 3**.

### *In vivo* screens read processing and count matrix generation

Base calls were demultiplexed with Cell Ranger 8.0.1 mkfastq and aligned to a custom reference comprising mouse GRCm38 (mm10) augmented with the RCAS proviral sequence as an additional contig. Count matrices were produced with cellranger count at default parameters. CRISPR Feature Barcode libraries were processed in parallel with a feature reference listing each library protospacer; the fixed mU6 *Cdkn2a*(ARF) guide carries no capture sequence and is therefore absent from this reference.

### gRNA assignment and quality control

Per-cell assignment used the Cell Ranger protospacer_calls_per_cell.csv output through a sequential pipeline applied identically to both screens: (i) multiplet rescue, collapsing cells with multiple guides to the dominant guide where the top-to-second UMI ratio exceeded 2.5; (ii) exclusion of cells retaining more than three guides; (iii) a minimum total guide UMI count of 3; (iv) collapse of the spacer suffix (_1-_5) to the target-gene level with within-cell de-duplication, preserving combinatorial labels (for example *Cdkn2a*|*Trp53*); and (v) a cohort-wide minimum of more than 13 cells per protospacer in the single screen and 5 in the dual screen, the lower dual cutoff chosen to preserve combinatorial arms diluted across a larger set of effective identities. Four non-targeting sequences constitute the Ntc set. Cells with no surviving assignment were excluded from perturbation-stratified analyses and retained for compartment-level and per-tumor analyses.

### *In vivo* screens single-cell preprocessing

The two screens were processed through independent pipelines from alignment onward to preserve per-mouse and per-screen heterogeneity. Doublets were removed with Scrublet per mouse at default parameters, and genes expressed in fewer than 5% of cells were filtered before downstream analysis. Dimensionality reduction used the top 80 principal components as input to mutual nearest neighbors batch alignment via Monocle3 align_cds with alignment_group = “batch” and residual_model_formula_str = “∼log_umi”, where batch denotes the 10x processing batch (four batches in the single screen and three in the dual). Clustering used Leiden community detection via cluster_cells(reduction_method = “Aligned”) at resolution 5 × 10⁻⁵, giving k = 10 clusters in each screen. UMAP was computed on the aligned space for visualization.

### Cell type annotation

Annotation proceeded in three stages. (i) Cells were scored against 16 canonical marker panels (tumor; microglia; monocyte-derived macrophage; border-associated macrophage; T cell; B cell; NK cell; neutrophil; mast cell; endothelial; pericyte; smooth muscle; fibroblast; oligodendrocyte; astrocyte; neuron; normal OPC) using an aggregate expression score, the natural log of one plus the sum of size-factor-normalized counts across each panel. (ii) Each cell was assigned the highest-scoring type provided the top score was ≥ 0.5 and the margin to the second ≥ 0.2; cells failing either were left unassigned. (iii) Each cell’s final label was set to the majority confident call within its Leiden cluster, provided ≥ 75% of confidently called cells carried the winning label and ≥ 10% of the cluster was confidently classified. Clusters whose majority fell below the winner threshold were not relabeled and their cells retained their own marker-based calls. A confidence-gated rescue additionally allowed a cell to retain its own call where it scored above the median of agreeing cells in its cluster. To separate tumor cells from parenchymal populations, clusters were designated tumor-confirmed where either the fraction of cells with a detected protospacer exceeded 5% or the fraction of RCAS-positive cells exceeded 50%.

### T cell subset annotation

Cells labeled T cell by the general cell type annotation procedure in either screen were subsetted and re-processed jointly. Leiden clusters were scored against marker panels for T-cell lineage (*Cd3d*, *Cd3e*, *Cd3g*, *Trac*, *Cd2*), coreceptor lineage (*Cd4*, *Cd8a*, *Cd8b1*), naive/memory (*Tcf7*, *Lef1*, *S1pr1*, *Bach2*, *Ccr7*, *Sell*, *Il7r*), Treg lineage (*Foxp3*, *Ikzf2*, *Ikzf4*, *Tnfrsf18*, *Nt5e*), activation (*Il2ra*, *Ctla4*, *Tnfrsf4*, *Tnfrsf9*, *Cd69*, *Cd44*), cytotoxicity (*Nkg7*, *Klrd1*, *Klrk1*, *Prf1*, *Gzmb*, *Ifng*) and γδ T cells (*Trdc*, *Zbtb16*, *Tmem176a*, *Tmem176b*, *Blk*, *Il17a*). Four states were assigned: naive T cell (n = 532), regulatory T cell (n = 301), γδ T cells (n = 46), and a population expressing the T-lineage panel without a dominant state program, reported as unpolarized T cells (n = 229).

### Human-to-mouse ortholog mapping

Published GBM signatures are defined on human symbols and were mapped to mouse symbols before scoring mouse data. Human symbols were first normalized for whitespace, including non-breaking spaces, before any pattern matching. Symbols were then resolved in a fixed order, most to least trustworthy: (i) direct match of the title-cased human symbol against the gene universe; (ii) ortholog lookup via babelgene, taking the best-supported mouse ortholog per human symbol; and (iii) a curated Mouse Genome Informatics (MGI)-based lookup for retired aliases and expanded families that babelgene does not resolve. The second and third steps accept a candidate only if that symbol is present in the gene universe. Symbols unresolved at this point were dropped rather than inferred. Where orthology is one-to-many because a human gene expanded into a mouse family, detection frequency in the tumor compartment was used to arbitrate among candidate members, preferring the member with materially higher detection and otherwise retaining the database pick.

### Verhaak/Wang bulk GBM subtype projection

Patchwork tumor cells and per-mouse pseudobulks were scored against the proneural, classical and mesenchymal programs of Verhaak et al. (2010) as refined by Wang et al. (2017), with the neural subtype retired. Pseudobulks were ortholog-mapped to human symbols and combined with TCGA-GBM bulk RNA-seq (log1p(CP10K)). Each signature was scored as a Tirosh-style aggregate module score (mean of signature genes minus the mean of an expression-bin-matched control set; 25 bins, 100 control genes per bin), and each unit assigned a subtype by argmax. Per-cell scores were computed identically and overlaid on the TCGA distributions (**Fig. 2c**).

### Neftel four-state per-cell classifier

Per-cell state assignment used the four-state meta-module framework of Neftel et al. (2019), with the classifier algorithm adapted from the framework of Nomura et al. (2025) and re-implemented in R, with one modification described below.

Each cell was scored against the six Neftel meta-modules (AC, OPC, MES1, MES2, NPC1, NPC2) and two cell-cycle modules using Tirosh background-bin-corrected module scoring. The combined state scores were then taken as the per-cell maximum across each two-component program. A per-program null was constructed by 20 rounds of gene-wise count shuffling within an Ntc-only reference pool and fitted to a normal distribution. For each cell, a one-sided upper-tail P-value was computed per state against the fitted null, the four P-values Holm-adjusted, and the highest-scoring significant program assigned (Padj < 0.05). Cells with no significant program were labeled unresolved. This yielded 24.8% resolved cells in the single screen and 27.3% in the dual. Restricting the null to unperturbed cells adapts the original framework to a perturbation screen, where pooling across all perturbations would inflate the null for the states most affected by perturbation. Hybrid detection and the two-dimensional quadrant projection (**Fig. 2d**, **3c**) followed Neftel et al. (2019) on the four-state score matrix.

### Per-perturbation Neftel state-shift inference

Perturbation-associated shifts in state score were assessed per tumor (**Fig. 2f**). For each state, a per-cell score was computed as the aggregate sum of log-normalized expression of that state’s signature genes and averaged within each tumor separately for focal and Ntc cells. Paired per-tumor differences were tested across tumors by two-sided Wilcoxon signed-rank test with Benjamini-Hochberg correction across the four states. Pairing within tumor controls for between-animal variation in baseline composition and treats tumor rather than cell as the unit of replication. The single-screen Ntc reference comprises 32 cells across eight tumors, so these comparisons are powered by the perturbed arms rather than by the reference.

### Neftel state calling in neurosphere cultures

Neurosphere cells were scored against the same six Neftel meta-modules with human symbols repaired against current HGNC nomenclature, using Monocle3 size-factor normalization (log₂(count/size factor + 1)) with Tirosh expression-matched control bins, and collapsed from six modules to four states by the per-cell maximum within each two-component program. States were assigned by argmax without a permutation null.

### Perturbation inclusion threshold

A per-perturbation inclusion threshold determined which perturbations were eligible for differential expression and other perturbation-stratified analyses. The threshold was set, independently within each screen, to the larger of the two internal-control populations recovered in that screen: the non-targeting controls, and *Mdm2*, whose loss is not expected to cooperate with *Pdgfb*-driven transformation. The rationale is that a perturbation represented by fewer cells than a control not expected to be selected cannot support a comparably powered effect-size estimate. In the single screen this is the *Mdm2* population (163 cells); in the dual screen it is the Ntc population (239 cells).

### Per-perturbation differential expression analysis

Differential expression was tested per perturbation in a focal-versus-other design. Focal cells were those whose guide label exactly matched the focal perturbation; comparators were all remaining guide-detected singlet cells in the same screen, including Ntc. Cells carrying combinatorial labels were excluded. A minimum of 30 focal cells was required, and the 5% expressed-gene filter was reapplied after subsetting. Models were fitted with glmmTMB using a negative-binomial parameterization with quadratic variance (nbinom2) and the formula counts ∼ arm + (1 | sample) + offset(log(size factor)), with the comparator as reference level and mouse as a random intercept. Non-converging fits were excluded. P-values were Benjamini-Hochberg corrected across tested genes within each perturbation. Volcano significance was defined as q < 0.05 together with a visualization-only effect-size cutoff of |β| > 0.10 (single) or |β| > 0.20 (dual), set at the inflection point of each screen’s β distribution. All inferential statements, gene-set definitions and correlation inputs use q < 0.05 alone.

### gRNA representation and enrichment

Representation of each target gene in tumors was compared against its representation in the injected DF1 pool, measured by nanopore amplicon sequencing of the plasmid preparation. Enrichment was computed per gene as log₂(fraction of guide-assigned tumor cells / fraction of DF1 amplicon reads), and tested by Fisher’s exact test with Benjamini-Hochberg correction across genes, at the cohort level and per tumor (**Fig. 2b**, **Fig. 4a**). Both sides of this comparison are expressed in target-gene space.

### Guide-level composition and clonality

Guide-level composition was reconstructed from protospacer calls retaining the spacer suffix, under the same UMI and enrichment criteria as the gene-level pipeline, restricted to QC-passing tumor cells (19,594 guide-resolved cells in the single screen and 14,591 in the dual, of which 17,287 and 11,282 respectively carried a single targeted gene). For each gene within each screen, the composition of its five protospacers was summarized as the proportion of guide-resolved cells carrying each, and compared against the DF1 pool.

Between-tumor heterogeneity was quantified as the mean pairwise Bray-Curtis dissimilarity between the five-element composition vectors of tumor pairs bearing that gene, restricted to genes with at least three tumors contributing ≥ 20 cells. Screens were compared per gene by a two-sided exact Wilcoxon signed-rank test over those paired genes.

### Gene-set enrichment analysis

Pre-ranked GSEA was performed with fgsea on the mixed-model differential expression output, ranking genes by sign(β) × −log₁₀(max(P, 10⁻³⁰⁰)) and collapsing duplicate symbols by averaging. Pathway P values were Benjamini-Hochberg adjusted within each run and significance set at P_adj < 0.05 unless stated otherwise.

### Navtemadlin signature derivation and comparison

The MDM2 inhibitor navtemadlin response signature was reconstructed from two published datasets of Rendo et al. (2025): (i) paired vehicle versus navtemadlin treatment of BT145 and BT286 *CDKN2A*-null, *TP53* wild-type human GBM patient-derived neurosphere cultures; and (ii) pre-treatment and 30 h on-treatment paired biopsies from 8 patients with recurrent *TP53* wild-type GBM. Per-line differential expression was computed with DESeq2 on the deposited normalized counts, and the two lines combined by a direction-aware Stouffer procedure. Only genes concordant in sign were retained, and the top 31 up and 94 down genes by within-direction meta-P selected, matching the published 125-gene signature and direction split. Ortholog mapping yields 96 genes shared with the per-screen tumor expression matrix.

### MrVI differential abundance and covariate-specific differential expression analysis

Sample-resolved latent structure was modeled with MrVI^35^, as implemented in scvi-tools v1.5.1. MrVI learns two latent representations per cell: a sample-unaware cell-state representation, u, and a sample-aware representation, z, obtained by composing u with a sample-specific effect learned by multi-head attention. The feature genes inputted to the model included the union of genes detected in more than 5% of cells across all cell types found in the Patchwork tumors. Model parameters were layer = “counts”, sample_key = “sample” (mouse-of-origin, 18 levels), batch_key = “10X_batch” (6 levels), n_latent_u = 10 and n_latent = 30.

Differential abundance across the dual- and single-screen was computed with sample_cov_keys = “screen”. For each cell, MrVI evaluates the log density of its *u* coordinate under each sample and aggregates these into a log probability per covariate level. The per-cell dual-versus-single log ratio was taken as the difference between the two, with positive values marking regions enriched for the dual-screen.

### Tumor-immune ligand-receptor analysis with MultiNicheNet

Cell-cell communication was inferred with MultiNicheNet, which combines multi-sample pseudobulk differential expression on sender and receiver populations with NicheNet ligand-activity inference and ranks interactions by an integrated prioritization score, using the mouse NicheNet v2 prior. Two analyses were run. In the cross-screen analysis (**Fig. 3g**), senders were the four Neftel-state tumor populations, the receiver was the microglia compartment, mouse was the sample identifier and screen the group identifier, contrasted as dual versus single. In the per-genotype analysis (**Fig. 4g,h**), tumor senders of each dual-screen genotype were contrasted against the pooled remaining genotypes, with T cells or microglia as receivers. Ligand differential expression was tested by edgeR v4.4.2 pseudobulk requiring at least 5 cells per pseudosample.

### Latent-variable perturbation modeling (SHERLOCK)

Perturbation effects were modeled in a shared latent space with SHERLOCK, an L0-gated variational autoencoder, fit separately to each screen on tumor cells from perturbations meeting the inclusion threshold. Input features were genes reaching significance in the per-perturbation mixed-model differential expression output: per perturbation, genes with q < 0.1, or the 200 lowest q-value genes where fewer than 200 met that cutoff, unioned across perturbations with mitochondrial and ribosomal genes removed and selected pathway genes added. Hyperparameters were chosen by grid search scored on a composite of gate bimodality, average treatment effect, and concordance with the differential expression effect sizes.

The single-screen model was fit with a single representative seed while the dual-screen model was fit under five random seeds. The dual screen latent embeddings and the perturbation grouping in **Fig. 4b,c** are shown for a single representative seed. The consensus matrix in **Fig. 4d** reports, for each perturbation pair, the fraction of seeds in which the pair fell in the same group under hierarchical clustering of the inferred perturbation covariance.

Per-perturbation gene weights were summarized as a signed mean shift. The SHERLOCK package computes target gene weights as mean_shift² / (var_total + 1e-8), where var_total is the log1p-CPM variance across control (Ntc) cells only, so |mean_shift| was recovered exactly as sqrt(EV × (var_total + 1e-8)). Magnitudes are the mean over five seeds. Each magnitude was signed by the corresponding per-perturbation DEG effect size β.

Target genes were selected by seed consensus: a gene was retained if, for at least one perturbation, it reached the model’s within-perturbation significance test at FDR < 0.05 in every seed. Hemoglobin, mitochondrial, ribosomal, heat-shock, vector and transgene sequences, unnamed numeric and Riken-clone identifiers, and sex-linked genes were excluded beforehand. Retained genes were clustered on the signed mean-shift matrix by average-linkage hierarchical clustering on correlation distance and cut into eleven clusters, incremented by one per additional 120 genes; gene order follows the dendrogram leaf order and perturbations are ordered by consensus group. Each target genes cluster was tested for over-representation of MSigDB Hallmark gene sets by hypergeometric test after mapping mouse symbols to human, using sets with at least five genes present and clusters with at least three mapped genes, with Benjamini-Hochberg correction across all cluster x set tests and a size-matched random-draw null alongside. The background is the model’s own feature universe.

**Fig. 4f** displays the interferon-response cluster across *Trp53*, *Pten*, *Rb1* and *Nf1*. The cluster was identified by content, as the one overlapping a fixed list of canonical interferon-stimulated genes most, rather than by cluster index, which is not stable across runs. Selection required all-seed significance in at least one of the four perturbations. Immune categories were assigned by a fixed precedence: a specific Reactome pathway where one applied, then MGI structural families, then Hallmark sets (interferon first), then MGI inducibility families, then broader Reactome parents, and finally a per-gene literature citation. Three sources were needed because the mouse Hallmark and Reactome collections are orthology-projected and return nothing for mouse-expanded interferon-effector families or H2 haplotype genes. Genes matching no category are shown as unassigned and not displayed.

## Data Availability

The single-cell RNA-sequencing data generated in this study will be deposited in the National Center for Biotechnology Information’s Gene Expression Omnibus and will be publicly available on publication. The *in vivo* screens comprise 18 tumor libraries (8 single-screen and 10 dual-screen), each with paired gene-expression and CRISPR Feature Barcode libraries, together with nanopore amplicon sequencing of the injected DF1 genomic DNA used as the input-composition baseline for both screens. All Patchwork plasmid constructs will be deposited at Addgene, and their whole-plasmid nanopore sequences will be available alongside them.

Previously published datasets reanalyzed in this study were obtained from public repositories: navtemadlin-treated GBM neurosphere RNA-sequencing (GEO GSE280891) and paired pre-treatment and on-treatment patient biopsies (GEO GSE280951) from Rendo et al. (2025); and TCGA-GBM bulk RNA-sequencing and clinical data from the Genomic Data Commons (portal.gdc.cancer.gov). Published gene signatures were obtained from Neftel et al. 2019 meta-modules, Verhaak et al. 2010 and Wang et al. 2017 bulk subtype signatures, and the navtemadlin response signature reconstructed from GSE280891 as described in Methods.

Source data for each main and supplementary figure panel are provided with this paper. Any additional information required to reanalyze the data reported here is available from the corresponding author on request.

## Acknowledgements

The authors would like to thank all members of the McFaline-Figueroa lab for helpful suggestions and critical discussion on the study. The authors also thank Erin Bush and the JP Sulzberger Columbia Genome Center for next-generation sequencing support, Wan-I Kuo of the Flow Cytometry Shared Resource for cell sorting support, and Yanping Sun of the Oncology Precision Therapeutics and Imaging Core for magnetic resonance imaging support.

## Author contributions

J.L.M.-F. conceived the project, analyzed and interpreted the data, and provided overall supervision of the study; N.H. and I.L. designed experiments; N.H. and I.L. analyzed and interpreted the data; N.H., I.L., R.M.G., B.V., S.P.A., M.V., J.G., D.W., T.N., L.S., V.L., and M.Z. performed experiments or aided in analysis. E.A. provided additional supervision. N.H., I.L. and J.L.M.-F. wrote the manuscript. All authors reviewed and contributed to the manuscript.

## Supplementary Figures

**Supplementary Figure 1.**
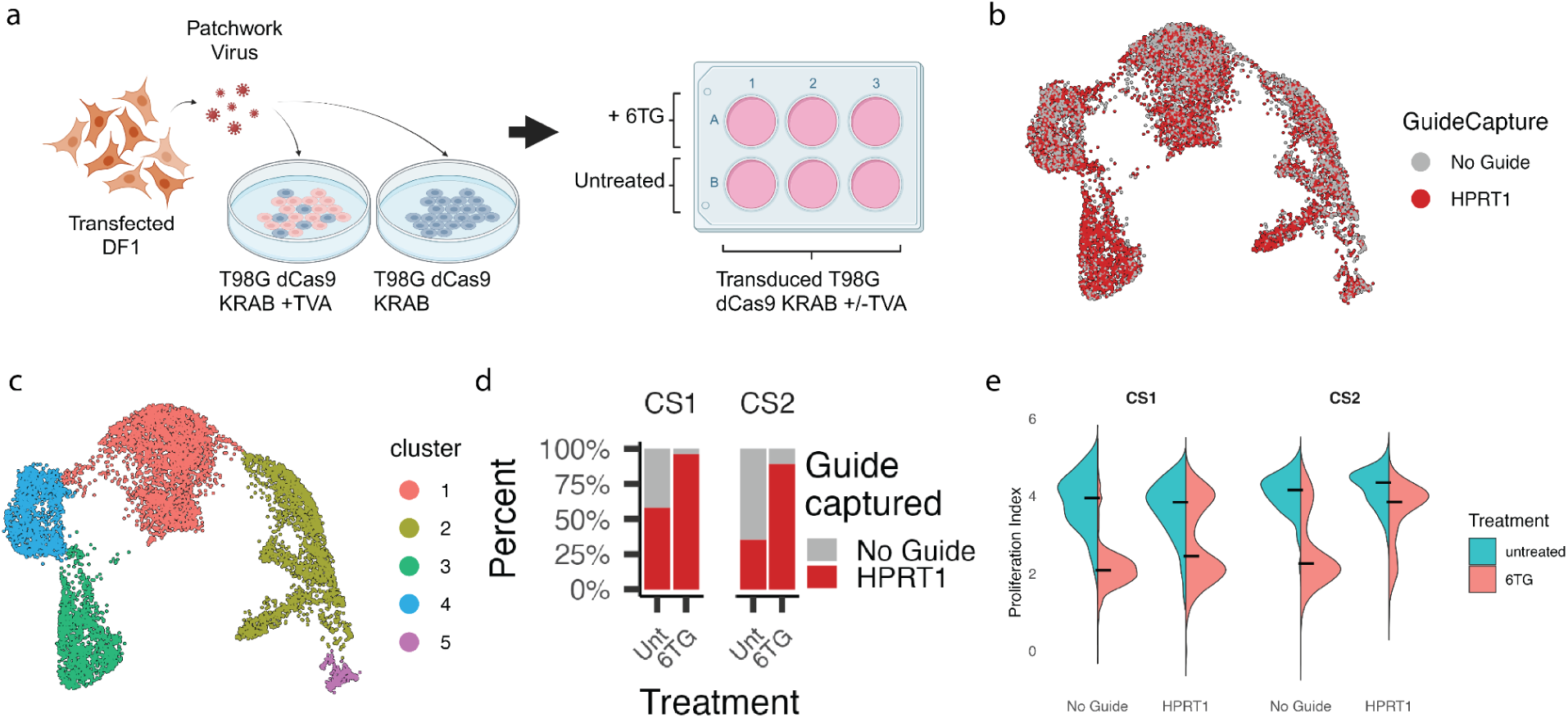
In vitro delivery of Patchwork avian viral particles enables gene editing in a cell-type-specific manner. **a,** Experimental schematic whereby T98G dCas9 KRAB (human GBM cells) expressing or not expressing receptor tvA are transduced with Patchwork retrovirus. The Patchwork viral vectors carry one of four constructs: RCAS CS1/2 sgHPRT1/Ntc. The transduced GBM cells were subsequently treated with 6-TG, and only GBM cells transduced with virus carrying sgHPRT1 were sequenced. Created in BioRender. McFaline-Figueroa, J.L. (2026) https://BioRender.com/e7491cn **b,** UMAP of the transduced T98G dCas9 KRAB + tvA mCherry cells annotated by gRNA captured, faceted by treatment condition and capture sequence. **c,** Same as B but annotated by Leiden clusters. **d,** Bar plots for the percentage of gRNAs captured within cluster 1 of the scRNA-seq data. Clusters were computed using the Leiden clustering algorithm. **e,** Violin plots of the proliferation index distribution across transduced T98G dCas9 KRAB + tvA mCherry cells in clusters 3 and 4, faceted by treatment condition and capture sequence.

**Supplementary Figure 2.**
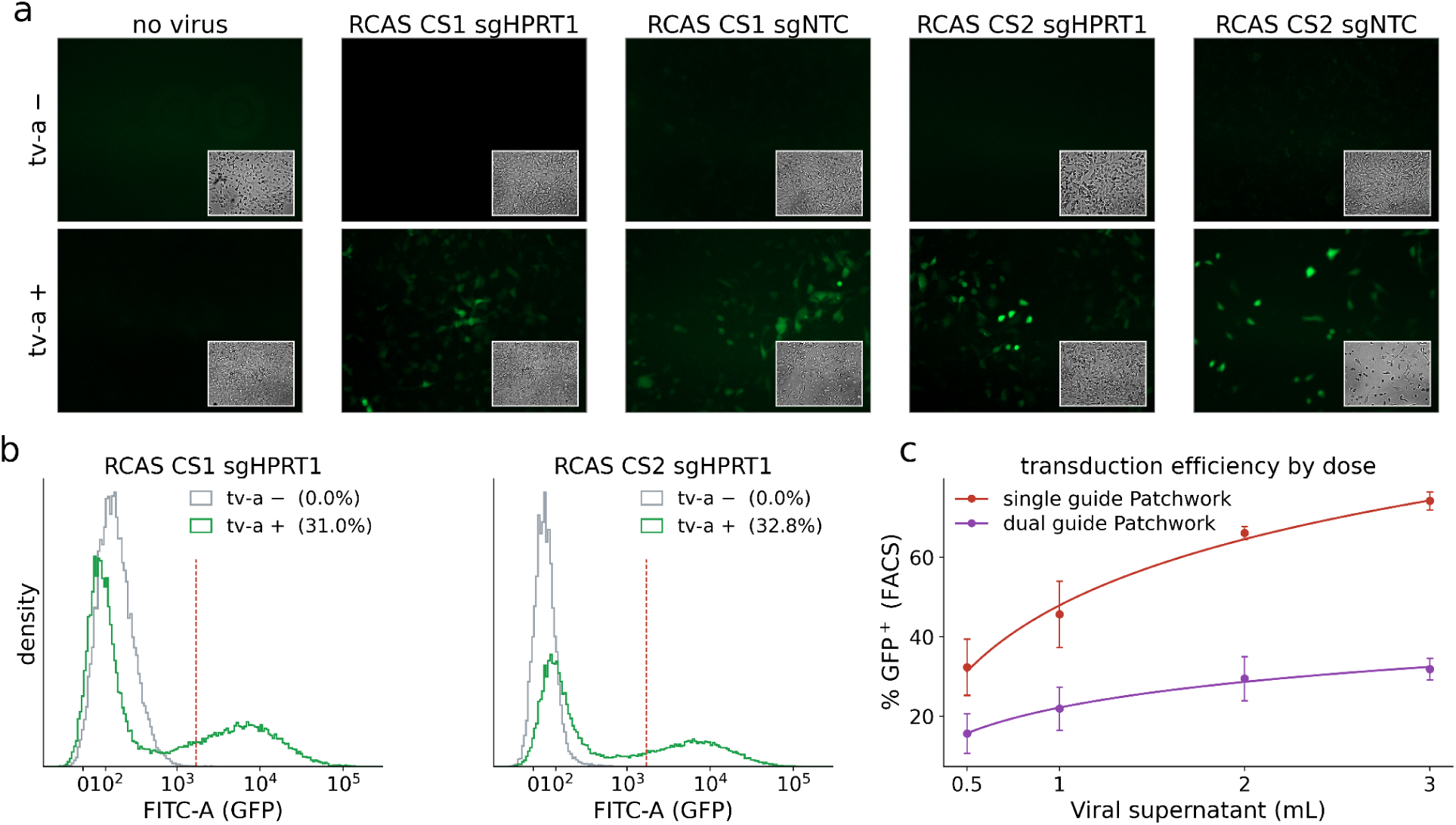
RCAS delivery of Patchwork constructs requires tvA and scales with construct design and viral dose. **a,** Fluorescence microscopy of human T98G dCas9 KRAB cells expressing or not expressing receptor tvA, exposed to Patchwork retrovirus carrying RCAS CS2 sgHPRT1 or to no virus. All GFP panels share one display range (22–150 of 255); brightfield insets confirm cell presence. Inset values give the percentage of image area above a GFP⁺ threshold of 40 of 255, set as the 99.99th percentile of the no-virus wells. **b,** Flow cytometric FITC-A (GFP) distributions for tvA-negative (gray) and tvA-expressing (green) cells transduced with RCAS CS1 or CS2 sgHPRT1. 50,000 events per sample, 92% retained after debris and singlet gating. The dashed line marks the FITC⁺ threshold of 1,700, the 99.9th percentile of the gated parental T98G reference; percentages of FITC⁺ cells are given in the legend. **c,** Percentage GFP⁺ in DF1 cells transduced with a range of Patchwork viral supernatant volumes containing either the single- or dual-guide Patchwork construct.

**Supplementary Figure 3.**
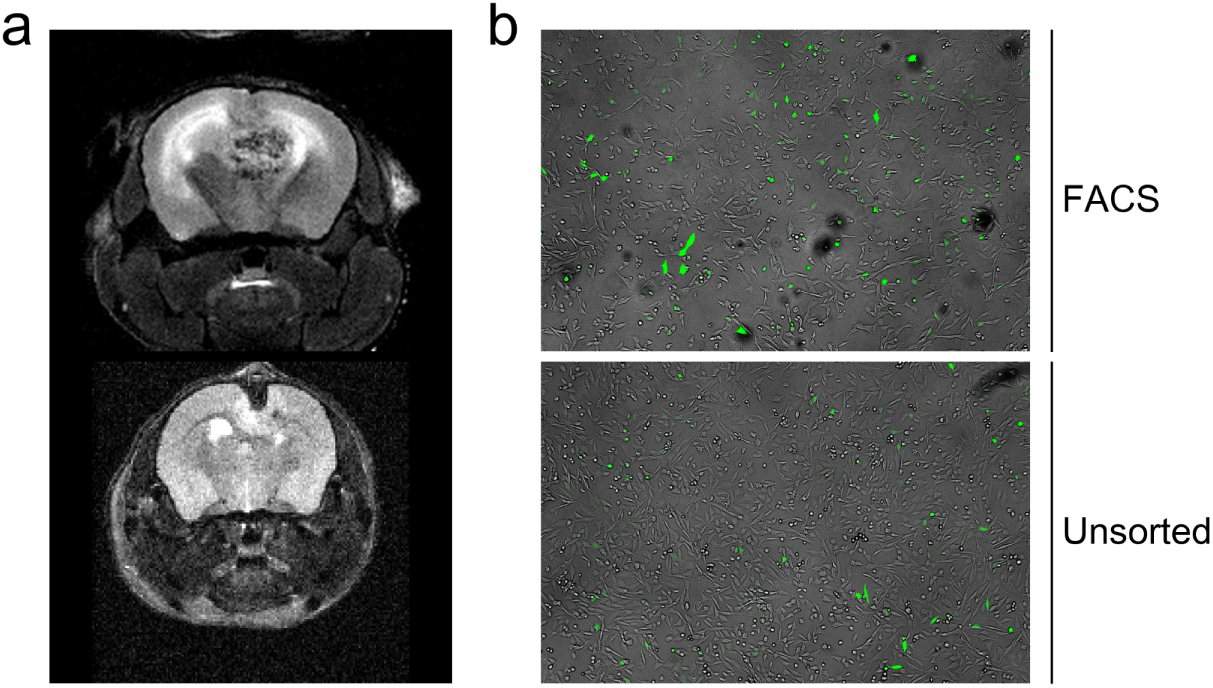
MRIs of Patchwork tumors in the single screen. **a,** T2-weighted coronal MRI of Nestin-tvA mice 25 days after intracranial injection of 1 × 10⁵ DF1 cells transfected with RCAS-CS2-sgTSG-PDGFB, either GFP⁺-sorted (top) or unsorted (bottom). **b,** Merged brightfield (grey) and GFP (green) microscopy images of the two transfected DF1 producer cells: sorted by FACS and unsorted.

**Supplementary Figure 4.**
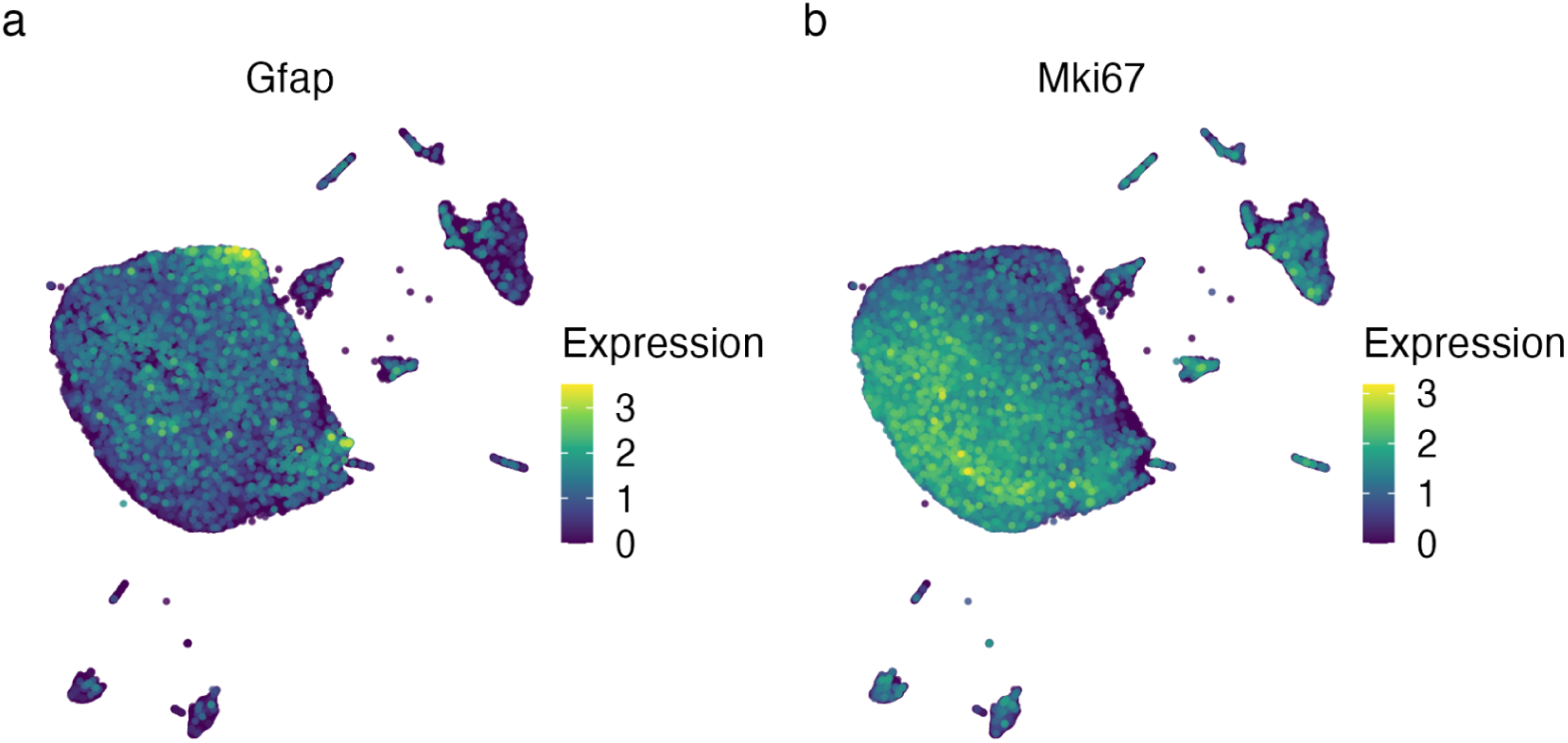
Single-screen UMAP annotated by *Gfap* and *Mki67* expression. UMAP embeddings of 121,514 cells from 8 mice in the single screen annotated by (**a**) *Gfap* expression and (**b**) *Mki67* expression.

**Supplementary Figure 5.**
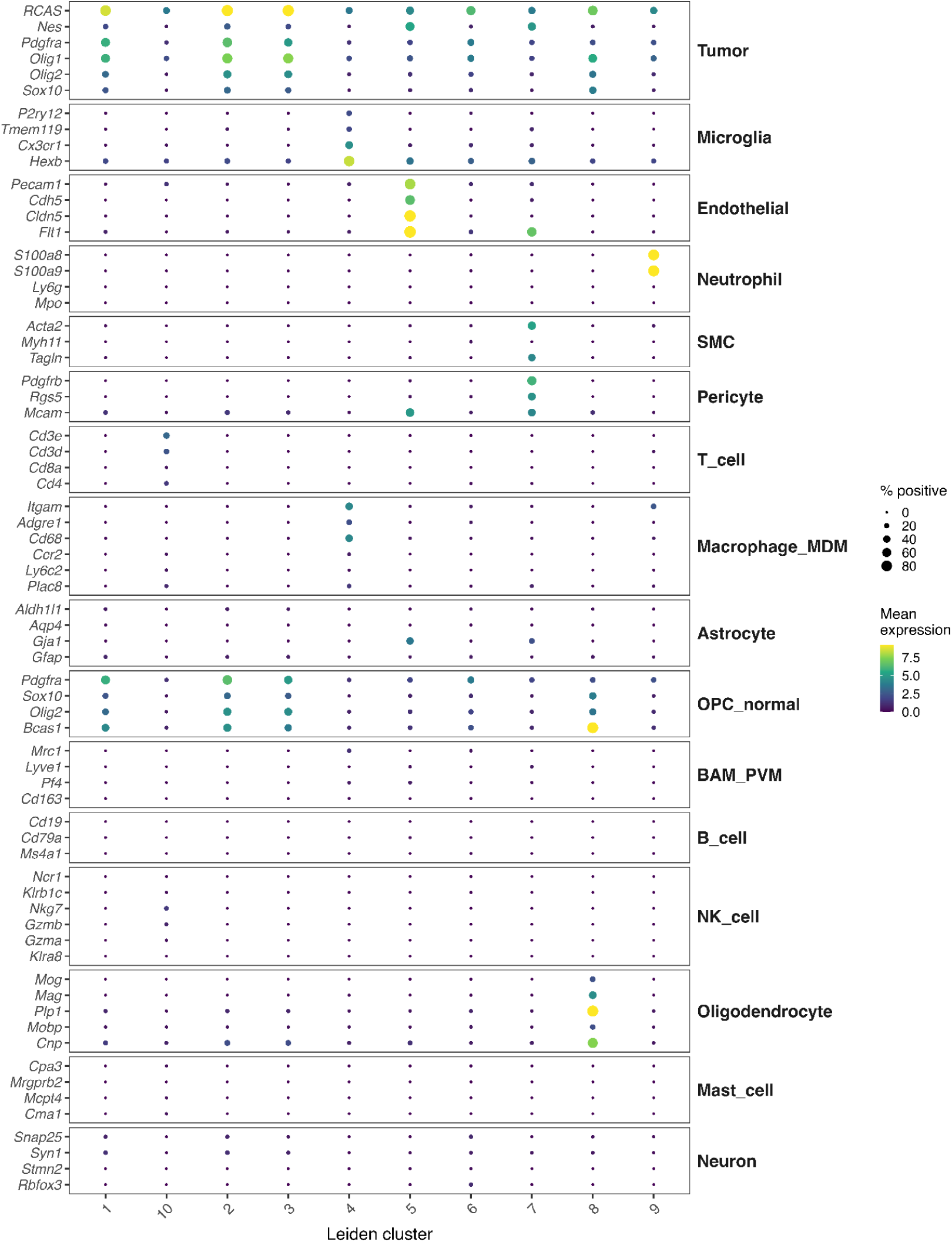
Cell type dot plot of cell types found in the single-screen. Per-cluster expression dot plot of canonical cell-type markers grouped by cell-type module. Dot size = % positive cells; color = mean expression per gene across clusters.

**Supplementary Figure 6.**
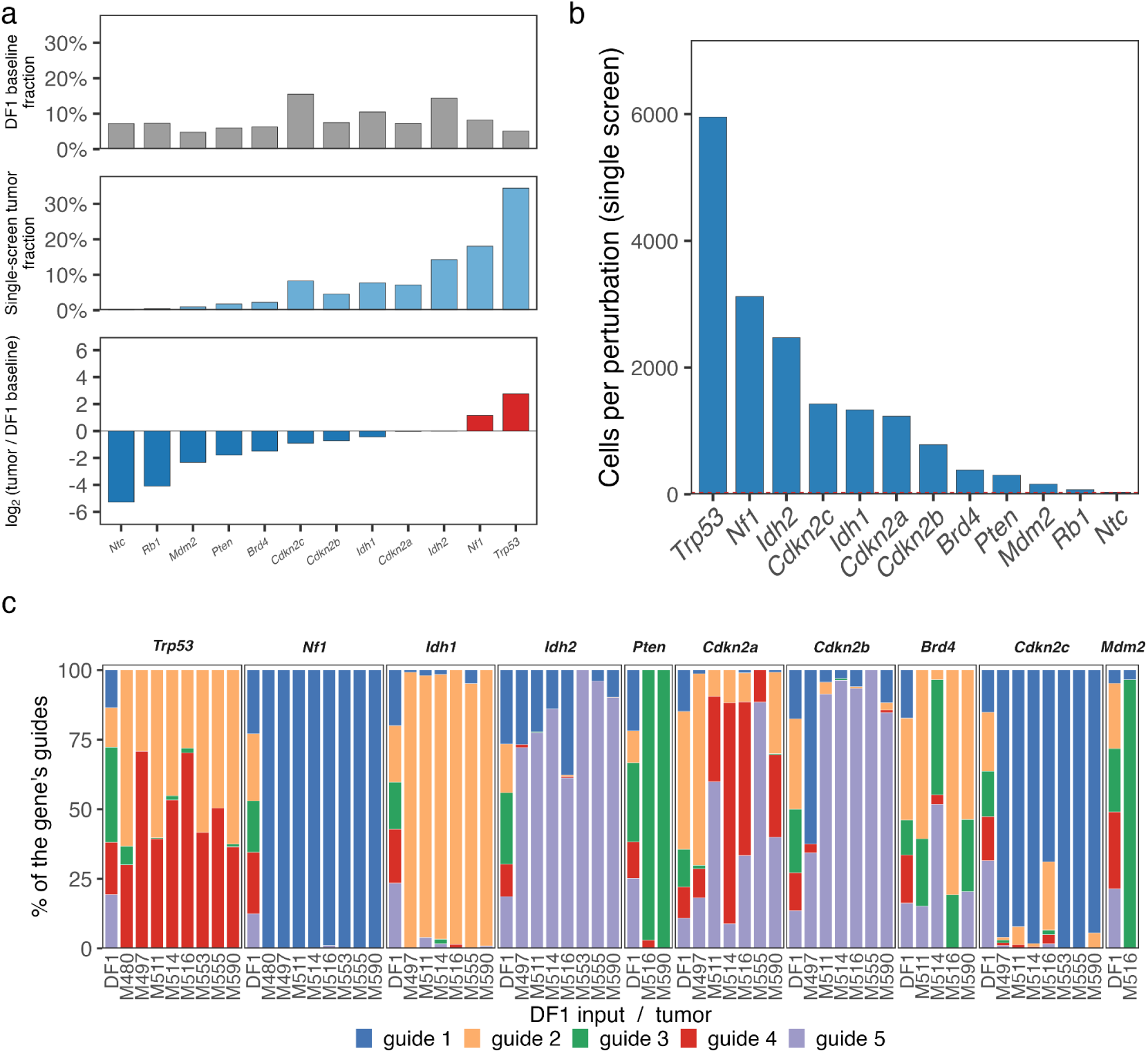
gRNA enrichment, per-perturbation representation, and guide-level composition in the single screen. **a,** Enrichment of each target gene from the injected library to tumor. Top: fraction of DF1 amplicon reads per gene in the baseline library (12 genes, including Ntc). Middle: fraction of single-screen tumor cells assigned to each gene. Bottom: per-gene log₂(tumor cell fraction / DF1 read fraction), colored red for enriched and blue for depleted, and sorted ascending. **b,** Cells recovered per perturbation in the single screen, sorted descending, with counts above each bar. **c,** Guide-level composition within each target gene, shown as the percentage of that gene’s cells carrying each of its five guides. Within each facet, the leftmost bar is the DF1 input library and the remaining bars are single-screen tumors. Genes are shown if at least 30 guide-resolved cells were recovered and individual tumors if at least 20 cells were recovered.

**Supplementary Figure 7.**
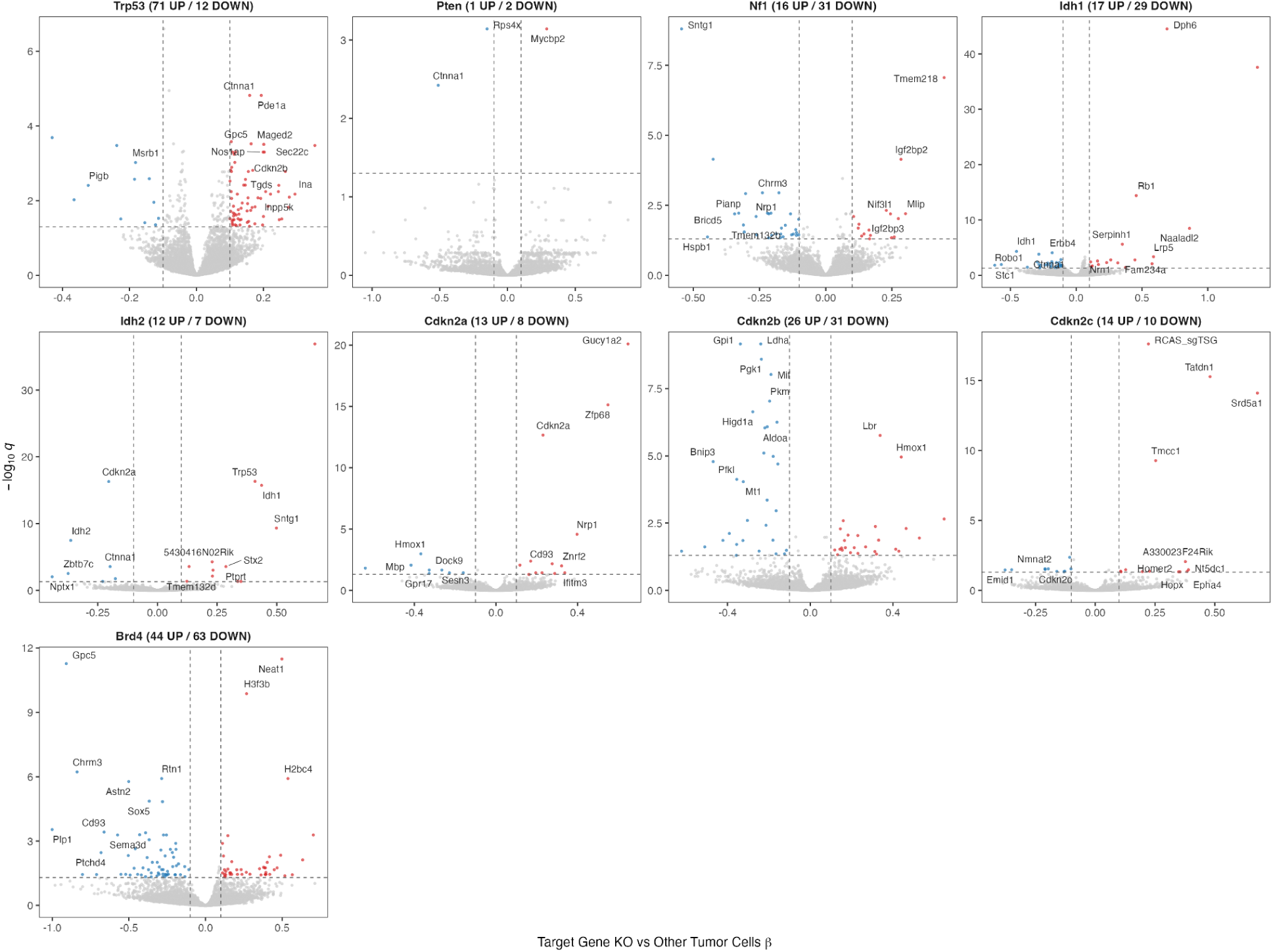
Per-perturbation differential expression in the single screen. DEG volcano per perturbation derived from a negative binomial GLMM comparing each perturbation to all other tumor cells and treating tumor as a random effect.

**Supplementary Figure 8.**
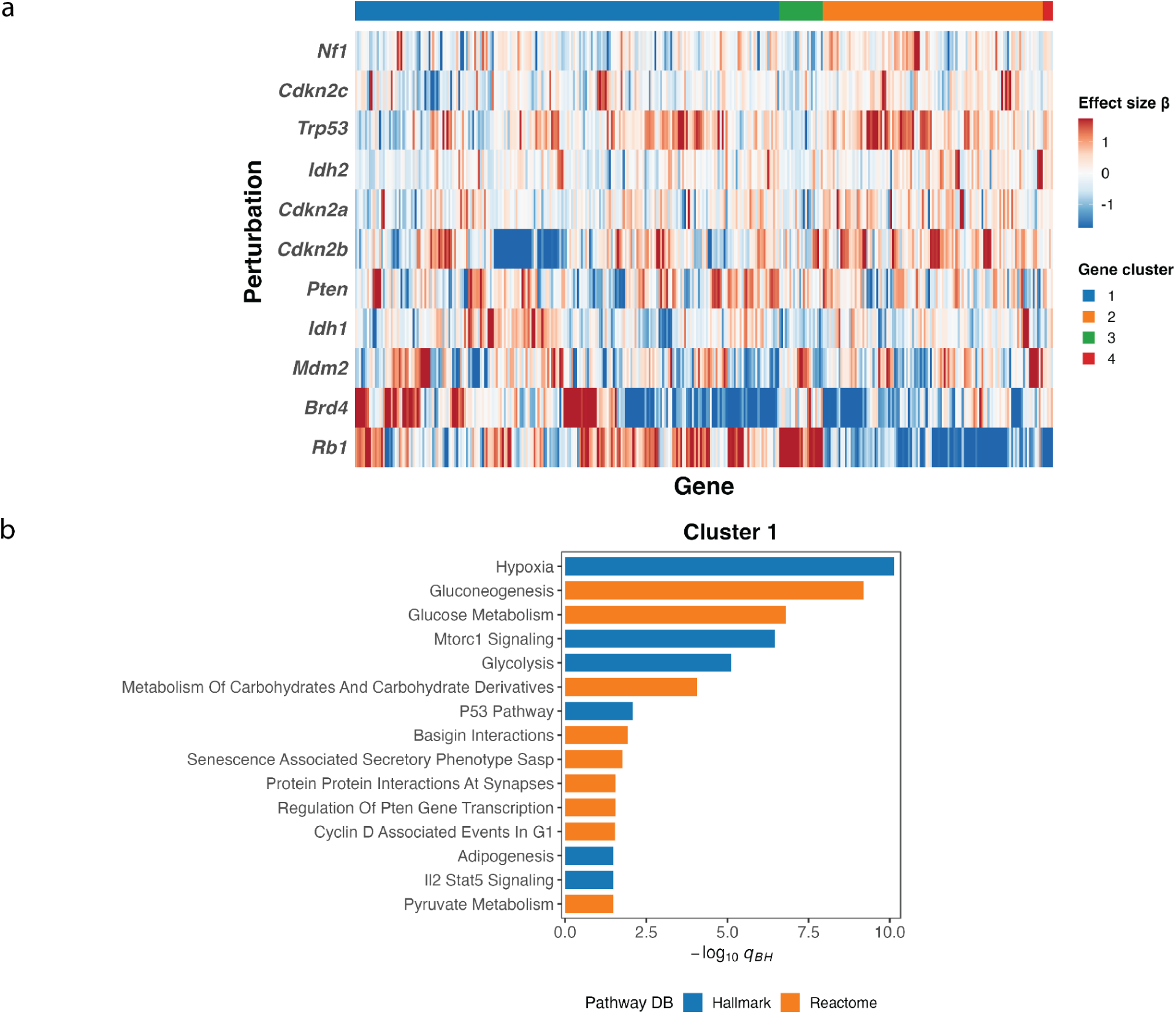
Single-screen per-perturbation DEG effect sizes heatmap and the derivative gene programs. **a,** Two-way hierarchical clustering of glmmTMB focal-vs-other effect sizes (β) across all eleven perturbations in the single screen. Genes were included if they reached q<0.05 (Benjamini–Hochberg) in at least one perturbation, giving 336 genes. Each gene’s β values were z-scored across perturbations and clipped at ±1.5 before plotting. Rows and columns were clustered on 1−Pearson correlation with Ward.D2 linkage and cut into four gene modules and four perturbation groups. **b,** GSEA testing of each gene module against MSigDB Hallmark and the Reactome gene sets by one-sided Fisher’s exact test, with Benjamini–Hochberg correction applied within each module. The background was the set of genes tested in the DEG models. Bars show the 15 most significant terms per module, ordered by odds ratio, for the three modules returning at least one term at q<0.05; the dashed line marks q=0.05.

**Supplementary Figure 9.**
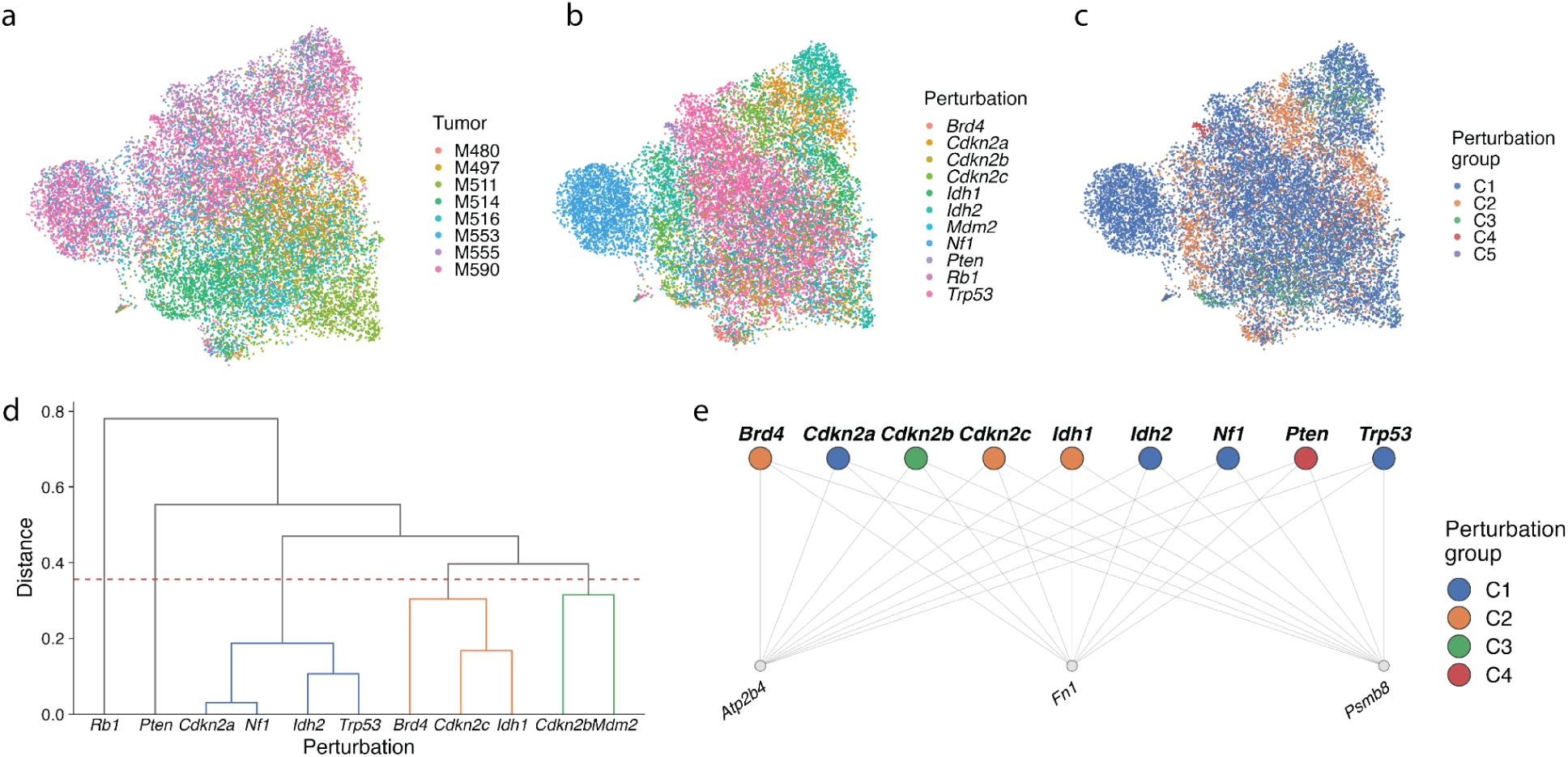
SHERLOCK latent-variable analysis of the single-screen. **a–c**, UMAP of the SHERLOCK per-cell latent representation of tumor cells with a non-Ntc guide assignment, annotated by tumor-of-origin (**a**), by the perturbation (**b**), and by the perturbation group (**c**). **d**, Hierarchical clustering of the learned perturbation covariance across the eleven perturbations. **e**, Bipartite graph of target genes shared by ten of the eleven perturbations; *Rb1* contributes none.

**Supplementary Figure 10.**
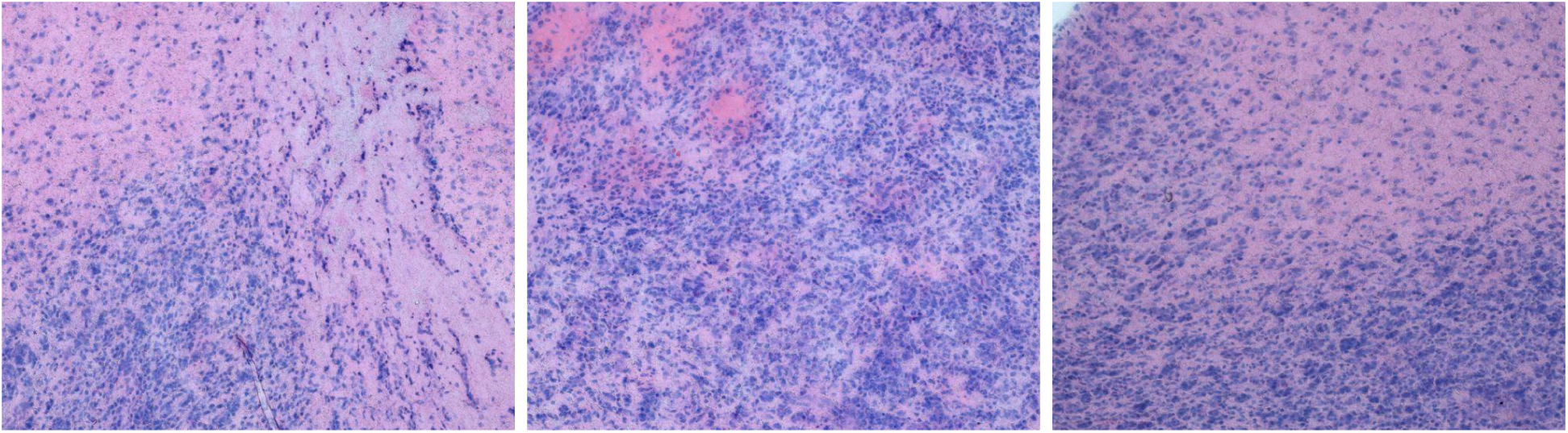
Histological H&E microscopy images of dual-screen tumors. H&E of a single representative Patchwork dual-screen tumor.

**Supplementary Figure 11.**
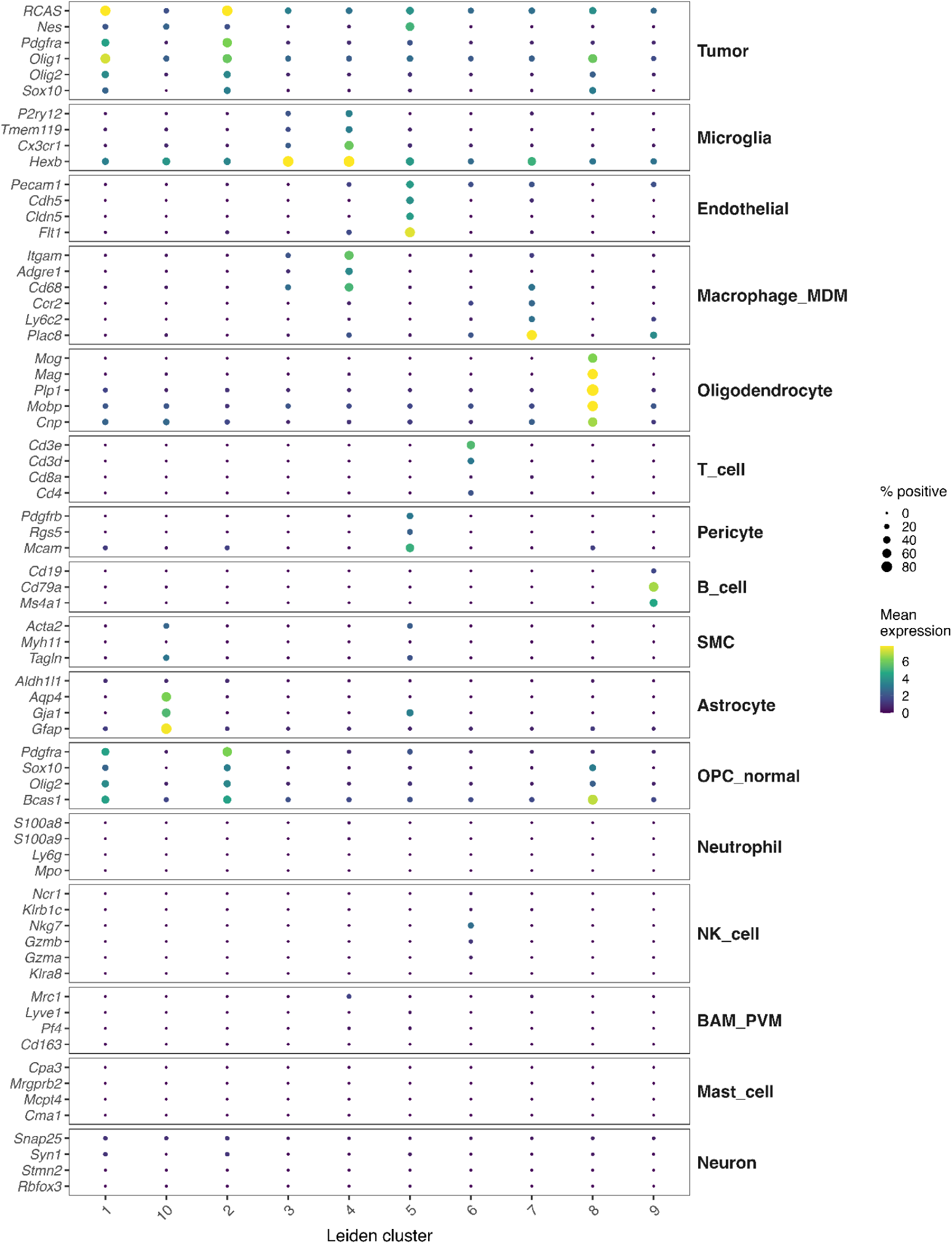
Cell type dot plot of cell types found in the dual screen. Per-cluster expression dot plot of canonical cell-type markers grouped by cell-type module. Dot size = % positive cells; color = mean expression per gene across clusters.

**Supplementary Figure 12.**
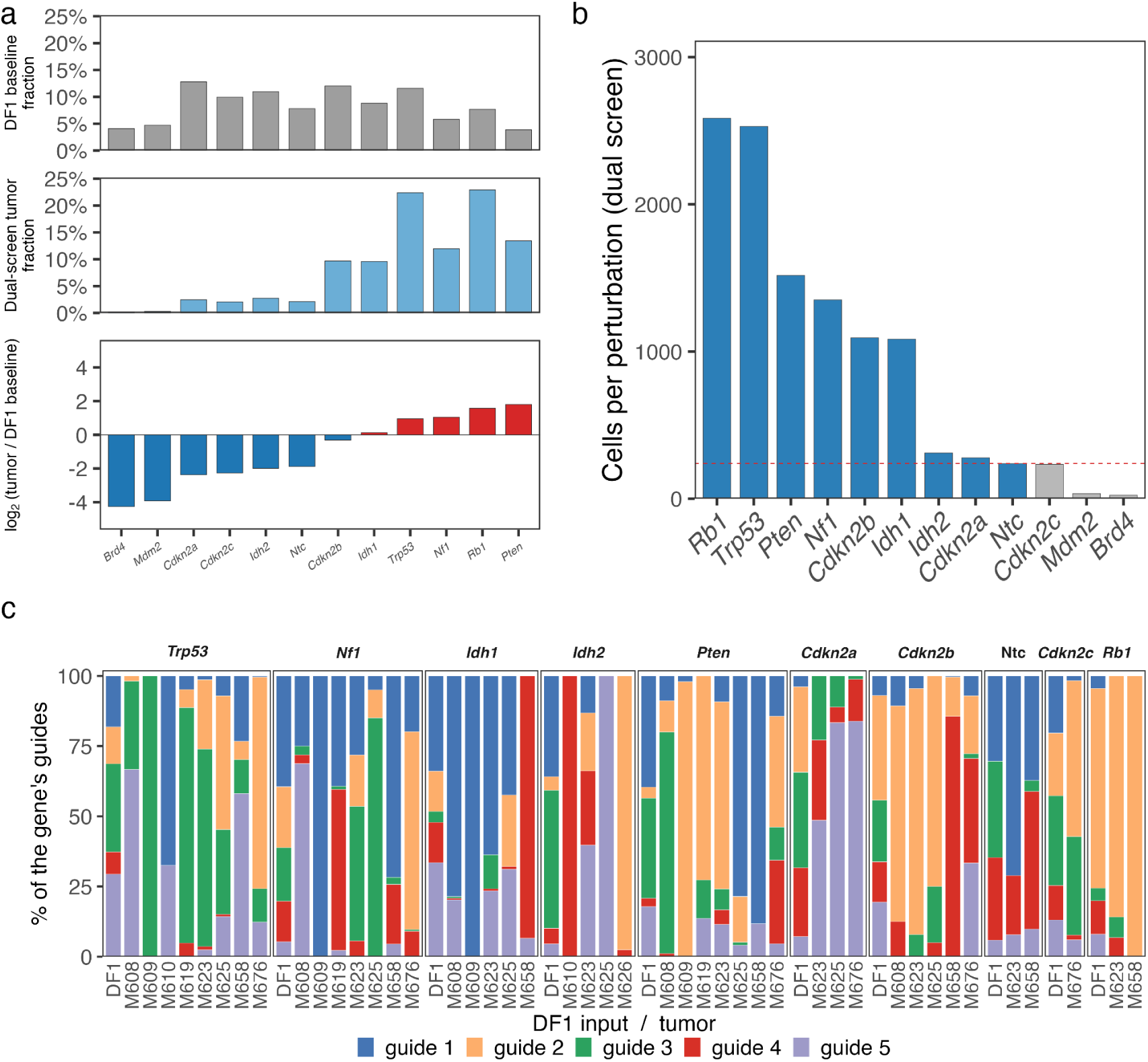
gRNA enrichment, per-perturbation representation and guide-level composition in the dual screen. **a,** Enrichment of each target gene from the injected library to tumor. Top: fraction of DF1 amplicon reads per gene in the dual screen baseline library (12 genes, including Ntc). Middle: fraction of dual screen tumor cells assigned to each gene. Bottom: per-gene log₂(tumor cell fraction / DF1 read fraction), colored red for enriched and blue for depleted, and sorted ascending. **b,** Cells recovered per perturbation in the dual-screen, sorted descending, with counts above each bar. The red dashed line marks the inclusion threshold for per-perturbation analyses, set to the screen’s total Ntc cell count (139 cells). **c,** Guide-level composition within each target gene, shown as the percentage of that gene’s cells carrying each of its five guides. Within each facet, the leftmost bar is the DF1 input library, and the remaining bars are dual-screen tumors. Genes are shown if at least 30 guide-resolved cells were recovered and individual tumors if at least 20 cells were recovered.

**Supplementary Figure 13.**
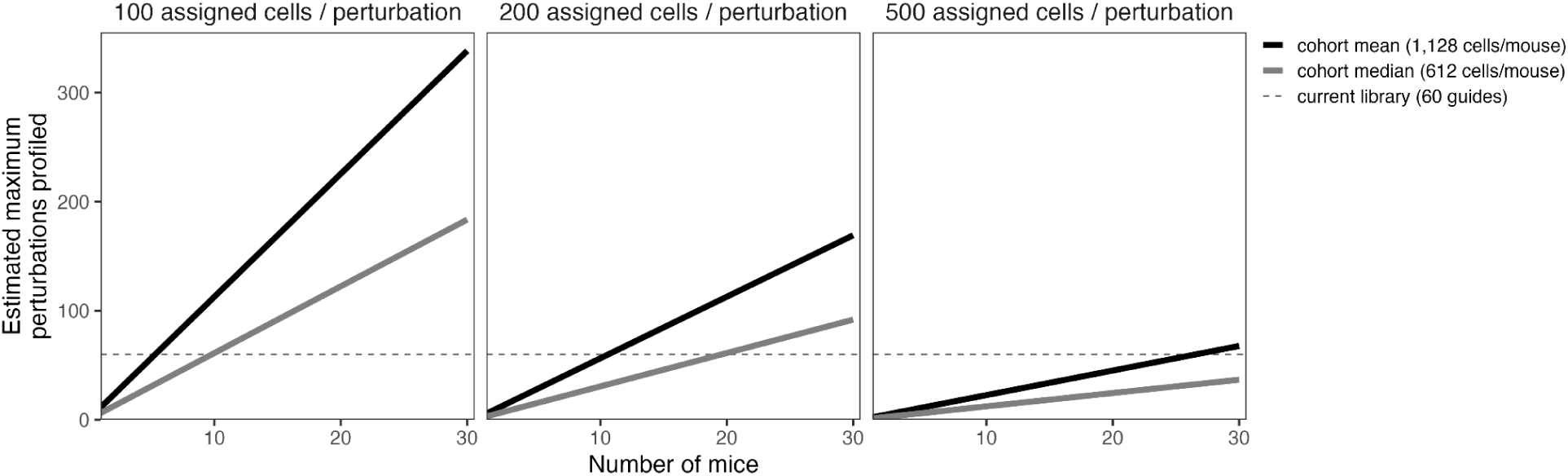
Estimated Patchwork library complexity with the dual-guide construct. Estimated maximum number of perturbations that can be profiled as a function of cohort size. Each panel fixes the number of assigned cells required per perturbation (100, 200 or 500); within a panel, perturbations supported = (mice × singleton cells per mouse) / cells per perturbation. The dashed line indicates the current tumor-suppressor library size (60 guides).

**Supplementary Figure 14.**
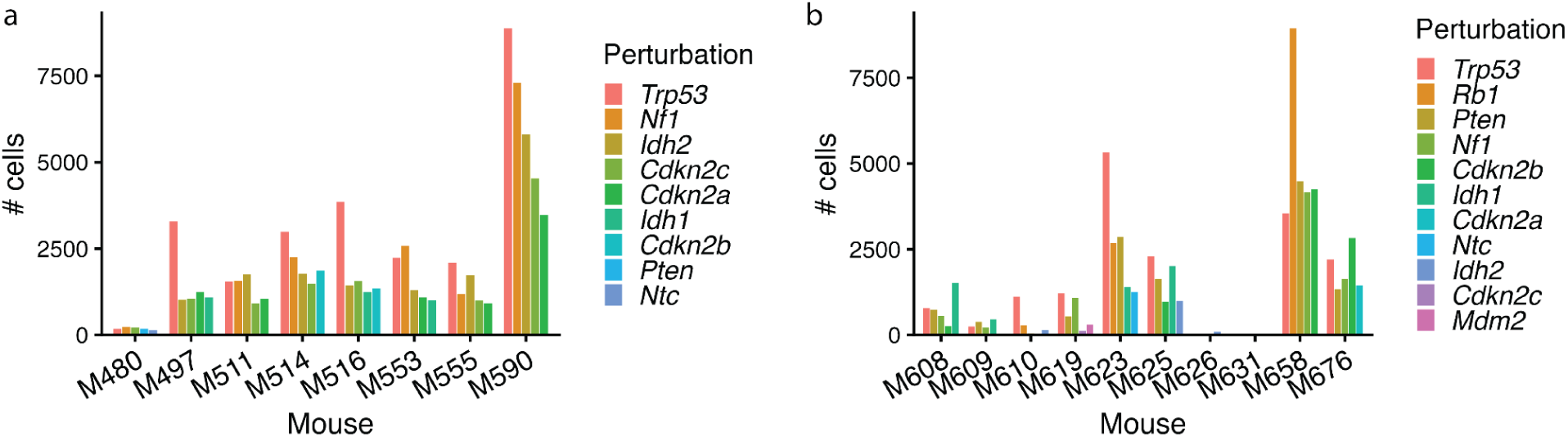
Per-tumor perturbation representation with a minimal guide capture criteria. **a,** Number of tumor cells carrying each perturbation in every mouse of the single-screen, calling a guide present whenever a cell carries ≥1 guide UMI for it. **b,** Same as (**a**) for the dual-screen

**Supplementary Figure 15.**
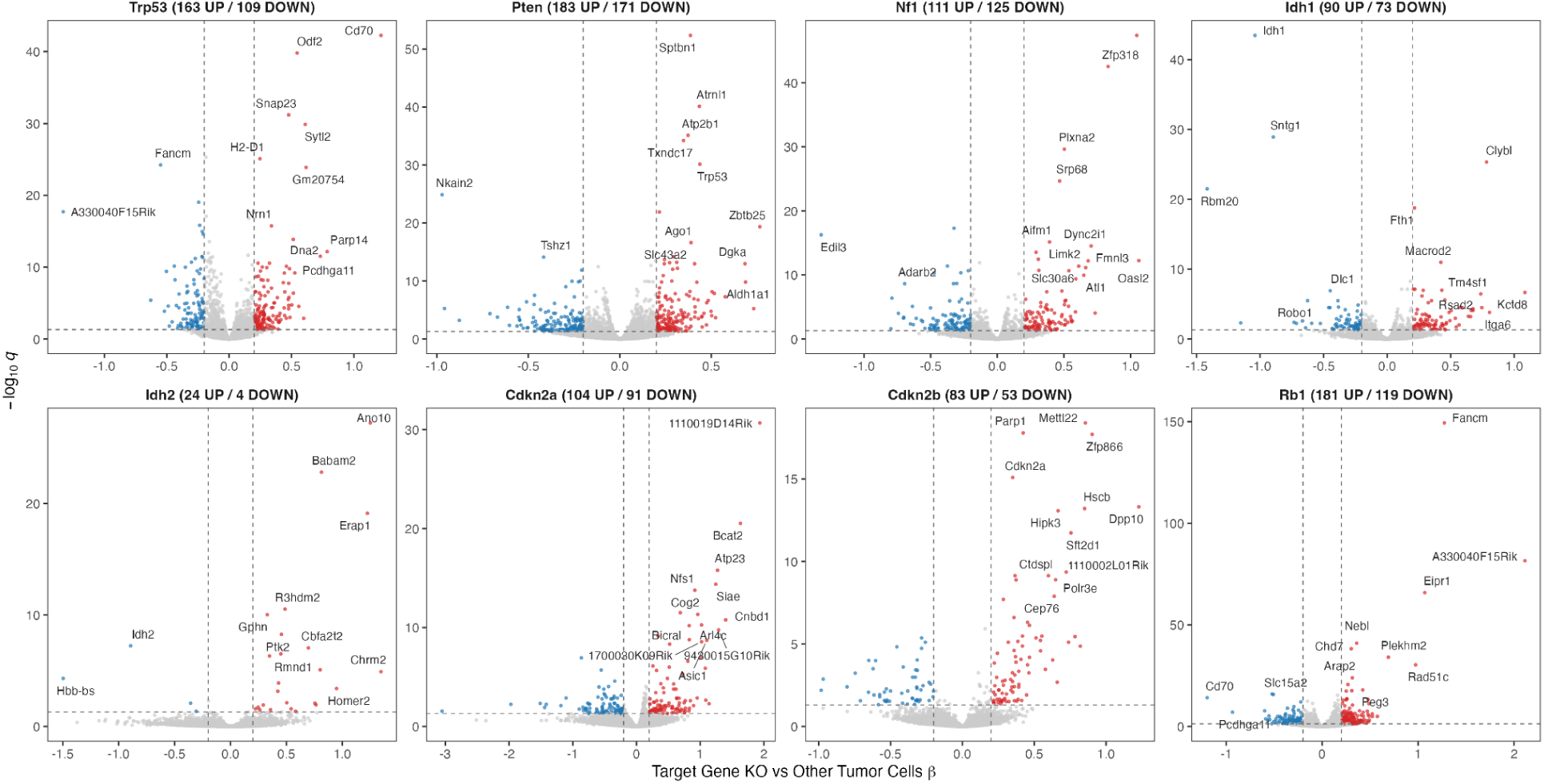
Per-perturbation differential expression in the dual-screen. DEG volcano per perturbation derived from a negative binomial GLMM comparing each perturbation to all other tumor cells and treating tumor as a random effect.

**Supplementary Figure 16.**
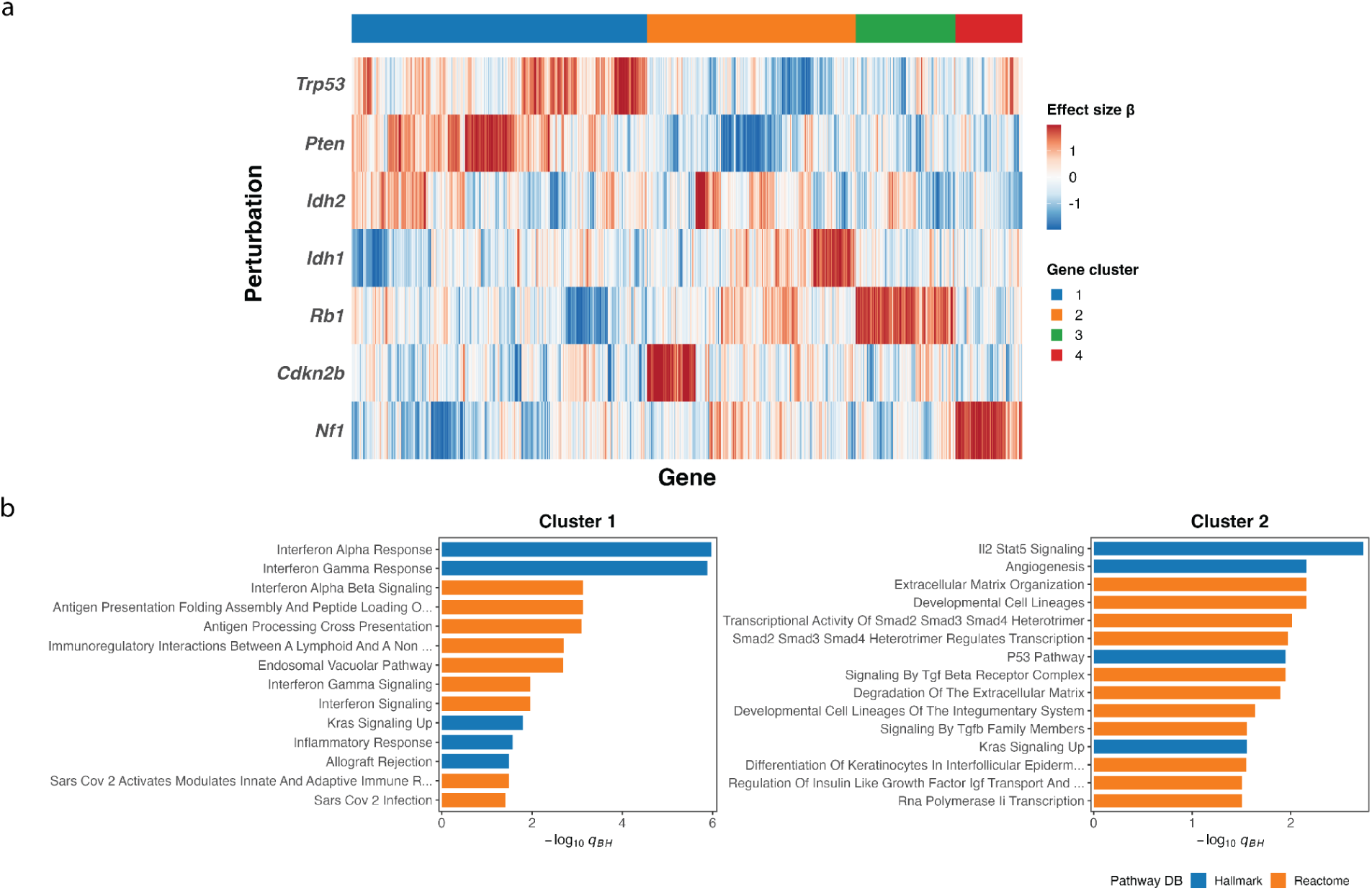
Dual screen per-perturbation DEG effect sizes heatmap and the derivative gene programs. **a,** Two-way hierarchical clustering of glmmTMB focal-vs-other effect sizes (β) across the seven perturbations of the dual-guide screen passing the inclusion criteria for per-perturbation analyses. Genes were included if they reached q<0.05 (Benjamini–Hochberg) in at least one perturbation, giving 1,137 genes. Each gene’s effect size β values were z-scored across perturbations and clipped at ±1.5 before plotting. Rows and columns were clustered on 1−Pearson correlation with Ward.D2 linkage and cut into four gene modules. **b,** GSEA testing of each gene module against MSigDB Hallmark and the Reactome gene sets by one-sided Fisher’s exact test, with Benjamini–Hochberg correction applied within each module. The background was the set of genes tested in the DEG models. Bars show the 15 most significant terms per module, ordered by odds ratio, for the three gene modules returning at least one term at q<0.05; the dashed line marks q=0.05.

**Supplementary Figure 17.**
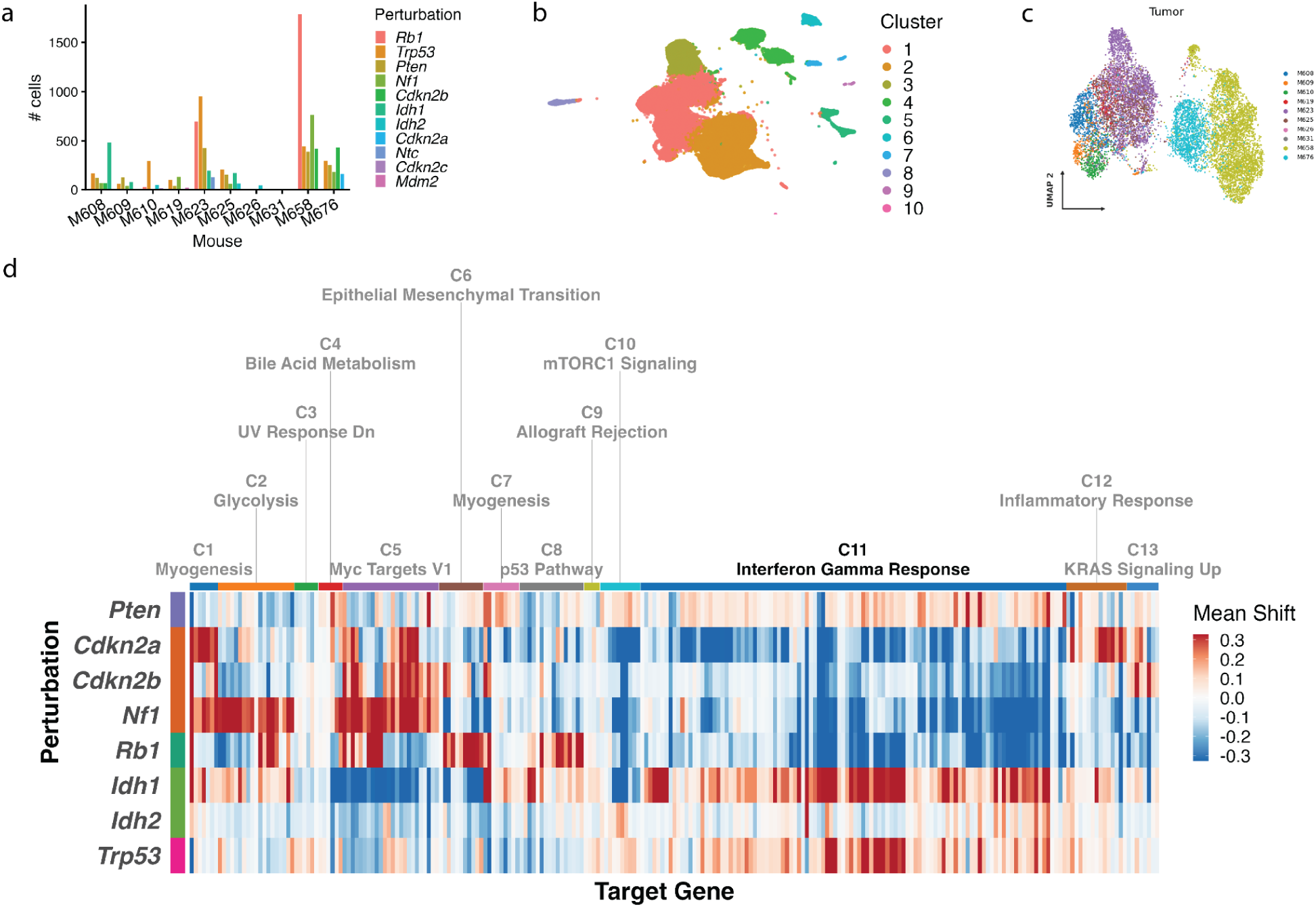
Perturbation-specific gene programs in the dual screen. **a**, Cells per perturbation in each dual-screen tumor. **b,** UMAP of the dual-screen cells annotated by Leiden cluster. **c**, UMAP of the SHERLOCK latent *z* space annotated by tumor, from a single representative seed. **d**, Signed mean shift per significant target gene across perturbations. The clustermap is annotated with nine unsupervised target gene clusters are, each labeled with the MSigDB Hallmark term most enriched in it by GSEA over-representation analysis.

**Supplementary Figure 18.**
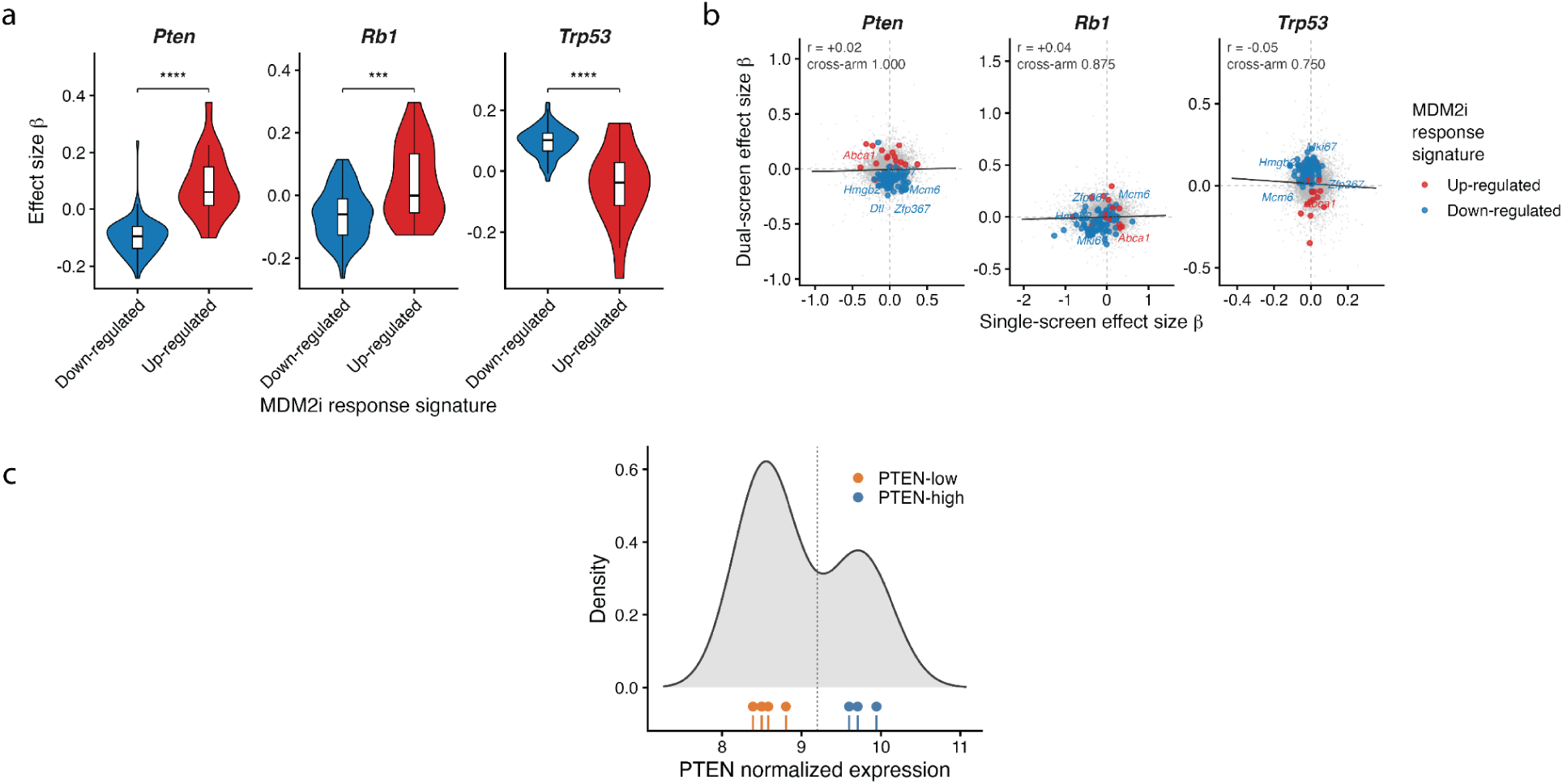
Pten and Rb1 loss lead to p53 hyperactivity resembling pharmacological MDM2 inhibition. **a**, Patchwork dual-screen DEG βs distribution comparing the up- and down-regulated arms of the navtemadlin MDM2i response signature across *Pten*, *Rb1* and *Trp53* KOs in the dual-screen. **b**, Patchwork dual-screen per-perturbation DEG effect size βs correlation against the matching single-screen βs, genome-wide over the 10,062 genes fitted in both screens. Navtemadlin response signature genes are highlighted. Insets give Pearson *r* with *r*², a gene-label permutation P-value, and a cross-arm P-value where cross-arm P indicates the fraction of the other perturbations whose single-screen β predicts this perturbation’s dual-screen β at least as well as its own does. **c,** Pre-treatment PTEN expression across the Rendo et al. 2025 patient cohort, with the dashed line marking the cut defining PTEN-low and PTEN-high. Ticks are individual patients.

**Supplementary Table 1. sgRNA library: protospacer sequences, design scores, screen representation and INK4 locus annotation.** One row per entry in the sgRNA library: 60 entries representing 59 distinct protospacers, comprising eleven targeted genes with five protospacers each plus five non-targeting controls. Library ID is the identifier used in the Cell Ranger feature reference and in the deposited sequencing data. Target gene is the gene the protospacer aligns to and is the key used throughout; for fifteen entries, it differs from the gene named in the Library ID. The design is described by protospacer, length, PAM, transcript, cut site on GRCm38 (mm10), exon, and cut position as a percentage of the coding sequence. On-target efficacy is the Rule Set 3 score, where higher indicates better predicted cutting. Aggregate CFD summarizes predicted genome-wide off-target burden, where lower is better and zero indicates that no off-target site was found. DF1 input reads are amplicon read counts from the transfected producer-cell pool, given separately for each screen. Tumor cells are the guide-resolved tumor cells recovered for that protospacer in each screen. The final columns annotate guides at the INK4 locus, where *Cdkn2a* exon 2 is shared between p16INK4a and p19ARF in different reading frames: locus, the transcript used for exon and cut position, reading frames at the cut, the protein products disrupted and their number, cut position against p16INK4a and against p19ARF, and a locus note. Note carries entry-specific remarks.

**Supplementary Table 2. Per-library sequencing quality control for the *in vitro* CRISPRi experiment.** Cell Ranger quality-control metrics for the four *in vitro* 10X libraries: CS1 and CS2 direct-capture constructs, each untreated or 6-thioguanine treated. Rows are transcriptome metrics (cells recovered, mean reads per cell, median genes and median UMIs per cell, total genes detected, total reads, valid barcodes, sequencing saturation, Q30 bases in the RNA read, reads confidently mapped to the transcriptome, and fraction of reads in cells) followed by CRISPR Feature Barcode metrics (mean reads per cell, sequencing saturation, fraction of reads usable, usable reads per cell, and the percentage of cells with at least one and with at least two guides detected).

**Supplementary Table 3.** Per-mouse sequencing depth, cell recovery, guide-assignment confidence, and animal metadata for the *in vivo* screens. One row per tumor for the eighteen *in vivo* mice, with a cohort row giving totals for counts and means or medians for depth metrics. Screen indicates the single-guide or dual-guide screen. Cells called by Cell Ranger and Cells retained give cell numbers before and after preprocessing. Cells with at least one guide UMI is the number of cells with at least one detected Feature Barcode read, with the corresponding percentage; guide-assigned cells is the number with a usable perturbation label, with its percentage, and serves as the denominator for per-perturbation analyses. Assigned cells fall into four confidence tiers that sum to the assigned total: confident, a single detected guide at five or more total guide UMIs; nominal, a single detected guide at three or four total guide UMIs; rescued, multiple detected guides in which the most abundant exceeds the second by more than 2.5-fold; and ambiguous, multiple detected guides retained under a combinatorial label, which therefore do not enter a single-gene perturbation arm. Distinct guide genes and distinct guide calls give, per tumor, the number of unique target genes and the number of unique labels including combinatorial ones. Median UMIs and median genes per cell, mean reads per cell for the gene-expression and CRISPR Feature Barcode libraries, and sequencing saturation are Cell Ranger per-library values. Time to harvest is measured in days from injection to collection.

